# G9a Promotes Breast Cancer Recurrence Through Repression of a Pro-inflammatory Program

**DOI:** 10.1101/2020.01.09.900183

**Authors:** Nathaniel W. Mabe, Shayna E. Wolery, Rachel Newcomb, Ryan C. Meingasner, Brittany A. Vilona, Chao-Chieh Lin, Ryan Lupo, Jen-Tsan Chi, James V. Alvarez

**Affiliations:** Department of Pharmacology and Cancer Biology, Duke University, Durham, NC 27710, USA; Department of Molecular Genetics and Microbiology, Duke University, Durham, NC 27710, USA

**Keywords:** G9a, breast cancer, recurrence, epigenetics, necroptosis

## Abstract

Epigenetic dysregulation is a common feature of cancer, and is thought to underlie many aspects of tumor progression. Using a genetically engineered mouse model of breast cancer recurrence, we show that recurrent mammary tumors undergo widespread epigenomic and transcriptional alterations, and acquire dependence on the G9a histone methyltransferase. Genetic ablation of G9a delays tumor recurrence, and pharmacologic inhibition of G9a slows the growth of recurrent tumors. Mechanistically, G9a activity is required to silence pro-inflammatory cytokines, including TNF, through H3K9 methylation at gene promoters. G9a inhibition induces re-expression of these cytokines, leading to p53 activation and necroptosis. Recurrent tumors upregulate receptor interacting protein kinase-3 (RIPK3) expression and are dependent upon RIPK3 activity. High RIPK3 expression renders recurrent tumors sensitive to necroptosis following G9a inhibition. These findings demonstrate that epigenetic rewiring – specifically G9a-mediated silencing of pro-necroptotic proteins – is a critical step in tumor recurrence and suggest that G9a is a targetable dependency in recurrent breast cancer.

## Introduction

Amplification of Human Epidermal Growth Factor Receptor – 2 (ERBB2/HER2) occurs in approximately 20% of human breast cancers (Howlader et al., 2014). Advances in HER2 targeted therapies have dramatically improved patient outcomes, however, one in four women will experience a tumor relapse (Haque et al., 2012). Of these tumor recurrences, 30-40% will lose HER2 amplification, rendering them insensitive to anti-HER2 therapies and resulting in lower patient survival (Hurley et al., 2006; Mittendorf et al., 2009). Thus, new strategies are required to overcome therapeutic resistance in HER2-discordant recurrent breast tumors. Tumor relapse is generally thought to result from selection for genetic mutations. As such, molecular profiling of recurrent breast tumors has largely focused on genomic alterations that promote HER2-independent resistance, such as loss of PTEN (Nagata et al., 2004), gain-of-function mutations in PIK3CA (Loibl et al., 2018), and amplification of MET (Shattuck et al., 2008).

However, it is increasingly appreciated that epigenetic dysregulation can also contribute directly to tumor relapse and therapeutic resistance (Brien et al., 2016; Sharma et al., 2010). In cell culture models, dependency on epigenetic reprogramming has been shown to induce rapid and reversible resistance to targeted therapies and cytotoxic therapies (Shaffer et al., 2017; Sharma et al., 2010). In human cancer models, epigenetic modulation through EZH2 mediates adaptive resistance to chemotherapy in non-small cell lung cancer patient-derived xenografts (Gardner et al., 2017). Patient data also support the role of epigenetic dysregulation in breast cancer recurrence. Global histone lysine hypoacetylation and DNA hypomethylation are associated with poor prognosis in breast cancer (Elsheikh et al., 2009; Selli et al., 2019; Suzuki et al., 2009), and transcriptional reprogramming is a hallmark of chemoresistant recurrent breast tumors (Yates et al., 2017). Consistent with these observations, the histone deacetylase (HDAC) inhibitor Entinostat prolongs survival in patients with recurrent breast cancer (Tomita et al., 2016). Together, these studies implicate epigenetic mechanisms in promoting drug resistance and breast tumor relapse. However, specific epigenetic alterations that underlie breast cancer recurrence and therapeutic resistance have not been well defined, and could identify clinically relevant targets in preventing or treating recurrent disease.

To gain insight into biological pathways driving tumor recurrence, we and others have used a genetically engineered mouse (GEM) mammary tumor model with conditional Her2 expression that mimics key features of breast cancer recurrence in women (Alvarez et al., 2013; Goel et al., 2016; Moody et al., 2002). Administration of doxycycline (dox) to MMTV-rtTA;TetO-Her2/neu (MTB;TAN) mice induces Her2 expression in mammary epithelial cells, leading to the formation of Her2-driven adenocarcinomas. Dox withdrawal leads to tumor regression, but a small population of tumor cells can survive Her2 downregulation and persist as minimal residual disease. After a latency of several months, these residual tumor cells spontaneously re-initiate proliferation and give rise to recurrent tumors. Importantly, these tumors recur independently of the Her2 oncogene, suggesting tumors have acquired Her2-independent bypass mechanisms for their growth, mirroring observations in HER2-discordant human breast cancers. Previous studies using HER2-driven recurrence models have identified Met amplification (Feng et al., 2014) and <u>Cdkn2a</u> deletions (Goel et al., 2016) as critical genetic drivers of tumor recurrence. While genetic alterations underlie some tumor relapses, not all tumors have a clear genomic basis for recurrence. We reasoned that a subset of recurrent tumors may leverage non-genetic mechanisms to adapt to and recur following HER2 withdrawal. Thus, characterizing epigenetic and transcriptional profiles of primary and recurrent tumors could identify non-genetic mechanisms by which tumor cells survive Her2 downregulation and form recurrent tumors. In the current study we used these GEM models to evaluate the contribution of epigenetic remodeling to breast cancer recurrence.

## Results

### Tumor recurrence is associated with widespread epigenetic remodeling

To gain insight into epigenetic changes associated with tumor recurrence, we derived cell lines from three primary and five recurrent tumors arising in MTB;TAN mice (Alvarez et al., 2013; Mabe et al., 2018). Prior work in several models has shown that Met amplification is a common genetic escape mechanism following oncogene withdrawal (Feng et al., 2014; Liu et al., 2011). We first characterized whether primary and recurrent tumor cells exhibit Met amplification. Copy number analysis revealed that while none of the primary tumor cells had Met amplification, 2/5 recurrent tumor cells had amplified Met between 5- and 15-fold **(Figure S1A)**. Met transcript levels followed the same pattern **(Figure S1B)**. We reasoned that in recurrent tumor cell lines #1-3, which lack Met amplification, tumor recurrence may instead be driven by epigenetic reprogramming.

To characterize epigenetic alterations in these tumors, we performed genome-wide ChIP-sequencing on primary cells (#1 and #2), and recurrent tumor cells (#1, #2 and #3). We evaluated histone marks H3K9ac and H3K4me3, which are commonly localized to actively transcribed genes, and the repressive histone mark H3K27me3, which is found at repressive heterochromatin (Wang et al., 2009). RNA polymerase II (RNApol2) was included to mark actively transcribed genes. Global analysis showed that each of the histone marks, as well as RNApol2, was enriched at the promoter and the transcription start site of active genes, as was expected based on their reported localization **(Figure S1C)**. To evaluate specific epigenetic alterations between primary and recurrent tumor cohorts, we performed differential binding (DiffBind) analysis for each histone modification and RNApol2. 29% of H3K4me3 peaks, 47% of H3K9ac peaks, and 18% of H3K27me3 peaks were differentially enriched in either primary or recurrent tumor cells, with recurrent cells having slightly more differential peaks than primary cells (Figure 1A). Half of the RNApol2 peaks we identified were differentially enriched in either primary or recurrent cells, with approximately equal numbers enriched in each cohort (Figure 1A). While most differential peaks were found at active gene regulatory elements, an increased number of peaks for H3K4me3, H3K9ac and H3K27me3 in recurrent cell lines were located in intergenic regions (>5 kb distance from a transcriptional start site; **Figure S1D)**. Intergenic localization of H3K9ac (Karmodiya et al., 2012), H3K4me3 (Pekowska et al., 2011) and RNApol2 (De Santa et al., 2010) has been described at enhancer or regulatory regions. To examine whether differential intergenic peaks overlapped with enhancers, we compared their localization with enhancer marks from an ENCODE dataset of mouse fibroblasts. There was a substantial overlap of differential H3K4me3 and H3K9ac intergenic peaks with the enhancer marks H3K4me1 and H3K27ac from ENCODE **(Figure S1E)**, suggesting that these intergenic peaks may represent newly acquired enhancers in recurrent tumor cells (Bradner et al., 2017).

**Figure 1:**
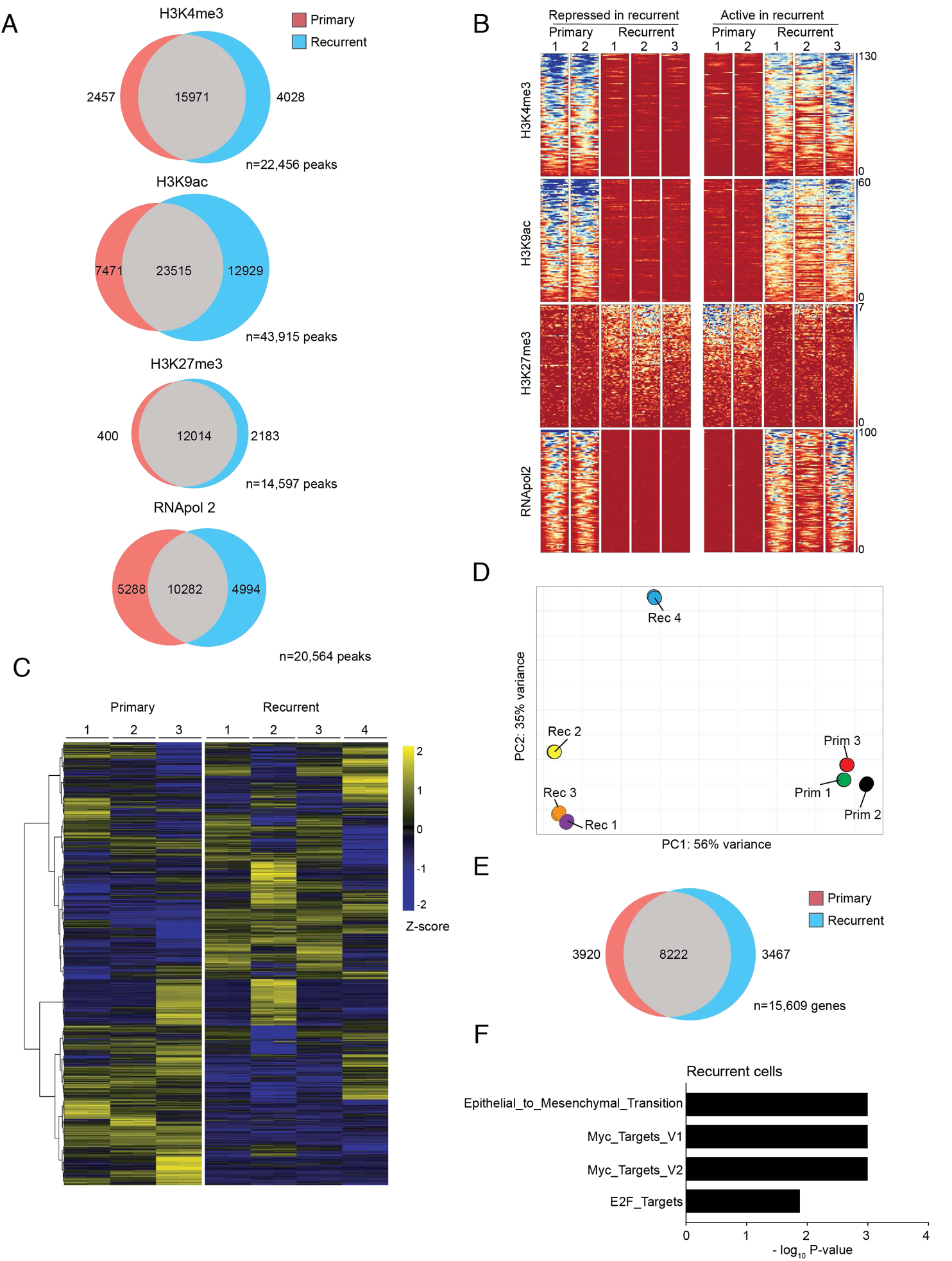
Tumor recurrence is associated with widespread epigenetic remodeling. A) Venn diagrams showing the number of ChIP-seq peaks unique to primary (n=2 cell lines) or recurrent (n=3 cell lines) tumors, and the total number of peaks analyzed for each epigenetic mark. B) Heatmap showing enrichment for active (H3K4me3 and H3K9ac) and repressive (H3K27me3) histone marks at the top 100 differentially RNApol2-bound genes in primary (left) and recurrent (right) tumor cell lines. Each row represents a different gene promoter. C) Heatmap showing unsupervised hierarchical clustering of 15,609 genes analyzed by RNA-sequencing for three primary and four recurrent tumor cell lines. Samples were performed in biological duplicate. Genes were median-centered and Z-score normalized within rows. D) Principal Components Analysis (PCA) of RNA sequencing from (C). E) Venn diagram showing the number of differentially expressed genes (adj. P-value < 0.05) between primary and recurrent tumor cell lines. F) Gene set enrichment analysis showing significantly enriched pathways in recurrent tumor cells.

We next compared the enrichment of these marks at individual gene promoters between primary and recurrent tumor cells. One group of genes was epigenetically repressed in recurrent tumor cells. The promoters of these genes had elevated enrichment of H3K9ac, H3K4me3, and RNApol2 and decreased enrichment of H3K27me3 in primary tumor cells compared to recurrent tumor cells (Figure 1B). Conversely, genes that were epigenetically activated in recurrent tumor cells displayed the opposite pattern, with elevated H3K9ac, H3K4me, and RNApol2 and decreased H3K27me3 peaks (Figure 1B). Taken together, these results indicate that tumor recurrence is associated with genome-wide epigenetic remodeling.

### Tumor recurrence is associated with transcriptional rewiring

Histone modifications are closely linked to the regulation of gene transcription. We reasoned that the genome-wide alterations in histone modifications between primary and recurrent tumor were likely associated with gene expression changes. To address this, we performed RNA-sequencing on three primary tumor cell lines (#1, #2 and #3) and four recurrent tumor cell lines (#1, #2, #3, and #4). Primary and recurrent tumors exhibited global differences in gene expression (Figure 1C) and clustered into distinct groups by principal components analysis (Figure 1D). Interestingly, the Met-amplified recurrent line (#4) had a gene expression pattern distinct from both primary and non-Met amplified recurrent tumor cells (Figure 1C **and** D), suggesting that Met amplification represents a distinct mode of tumor recurrence. Differential gene expression analysis identified 3,467 genes upregulated in recurrent tumor cells, and 3,920 genes upregulated in primary tumor cells (P-value adj. <0.05) (Figure 1E). These gene expression changes were strongly correlated with alterations in histone modifications at the gene promoters **(Figure S1F)**. Taken together, these results show that recurrent tumors exhibit genome-wide transcriptional changes that are associated with epigenetic rewiring.

To identify specific transcriptional programs altered in recurrent tumors, we performed Gene Set Enrichment Analysis (GSEA). Recurrent tumors were highly enriched for an epithelial-to-mesenchymal transition (EMT), as we and others have previously described (Mabe et al., 2018; Moody et al., 2005), as well as two independent Myc signatures (Figure 1F). Interestingly, a recent study in triple-negative breast cancer reported that EMT and Myc signatures are enriched in drug-resistant breast cancers (Kim et al., 2018). A number of genes found in the chemoresistant gene set were also upregulated in recurrent tumors, including Col1a1, Myc, Psat1, Cox6c, and Gastp1.

Further analysis of ChIP-seq data revealed that a substantial proportion of genes in both primary and recurrent tumor cells had bivalent promoters, marked by both the repressive histone mark H3K27me3 and the active histone mark H3K4me3 **(Figure S1G)**. Alterations in the expression of genes with bivalent promoters has been implicated in chemoresistance and enhanced breast cancer tumorigenicity (Chaffer et al., 2013; Chapman-Rothe et al., 2013). Consistent with this, we found a number of genes that were silenced through bivalent histone modifications in primary tumors and transcriptionally activated in recurrent tumors, including Mycn, Prrx2, Twist2, and Zeb1. Conversely, we identified two pro-apoptotic proteins, Bik and Pawr, that were expressed in primary tumors and silenced by bivalent regulation in recurrent tumors **(Figure S1G)**. Taken together, combined RNA- and ChIP-sequencing analyses indicate that tumor recurrence is associated with extensive epigenetic and transcriptional alterations.

### Recurrent Tumor Cells are Dependent on G9a Methyltransferase Activity

In light of the reported links between epigenetic remodeling and drug resistance (Knoechel et al., 2014; Sharma et al., 2010), we hypothesized that epigenetic remodeling may be functionally important for the survival and recurrence of tumor cells following oncogene withdrawal. Specifically, we reasoned that tumor cells that persist following oncogene withdrawal may acquire unique epigenetic dependencies that are not present in the primary tumors. To explore this hypothesis, we carried out a small-molecule screen testing the effect of inhibitors targeting various epigenetic enzymes on primary and recurrent tumor cell viability **(Table S1)**. Two primary (primary #1 and primary #2) and two recurrent (recurrent #1 and recurrent #2) tumor cell lines were treated with increasing doses of each inhibitor and cell viability was measured (Figure 2A). To identify drugs that differentially inhibit the growth of primary and recurrent tumor cells, we calculated the difference in IC_50_ for primary and recurrent tumor cells for each drug; values less than 0 correspond to drugs more potent against recurrent tumor cells. This analysis identified two drugs that were significantly more potent in recurrent cells, the G9a inhibitor BIX-01294 (IC_50_ difference=-0.88 [95% CI, −1.23 - −0.543]) and the Aurora kinase inhibitor lestaurtinib (IC_50_ difference= −1.06 [95% CI, −1.47 - −0.665) (Figure 2B). No drugs were significantly more potent against primary tumor cells (Figure 2B). Because we could not calculate an IC_50_ for nine compounds, we also evaluated the difference in maximal drug efficacy for each inhibitor between primary and recurrent cells (Figure 2C). Six drugs were significantly more efficacious in recurrent tumor cell lines (adj. *P*-value <0.05), including a G9a inhibitor (BIX-01294), a JMJD3 inhibitor (GSKJ1), an EZH2 inhibitor (EPZ5687), a DOT1L inhibitor (SGC0946), an LSD1 inhibitor (GSKLSD1), and a BAZ2A/B inhibitor (GSK2801) (Figure 2C). No inhibitors were significantly more efficacious in primary tumor cell lines (Figure 2C).

**Figure 2:**
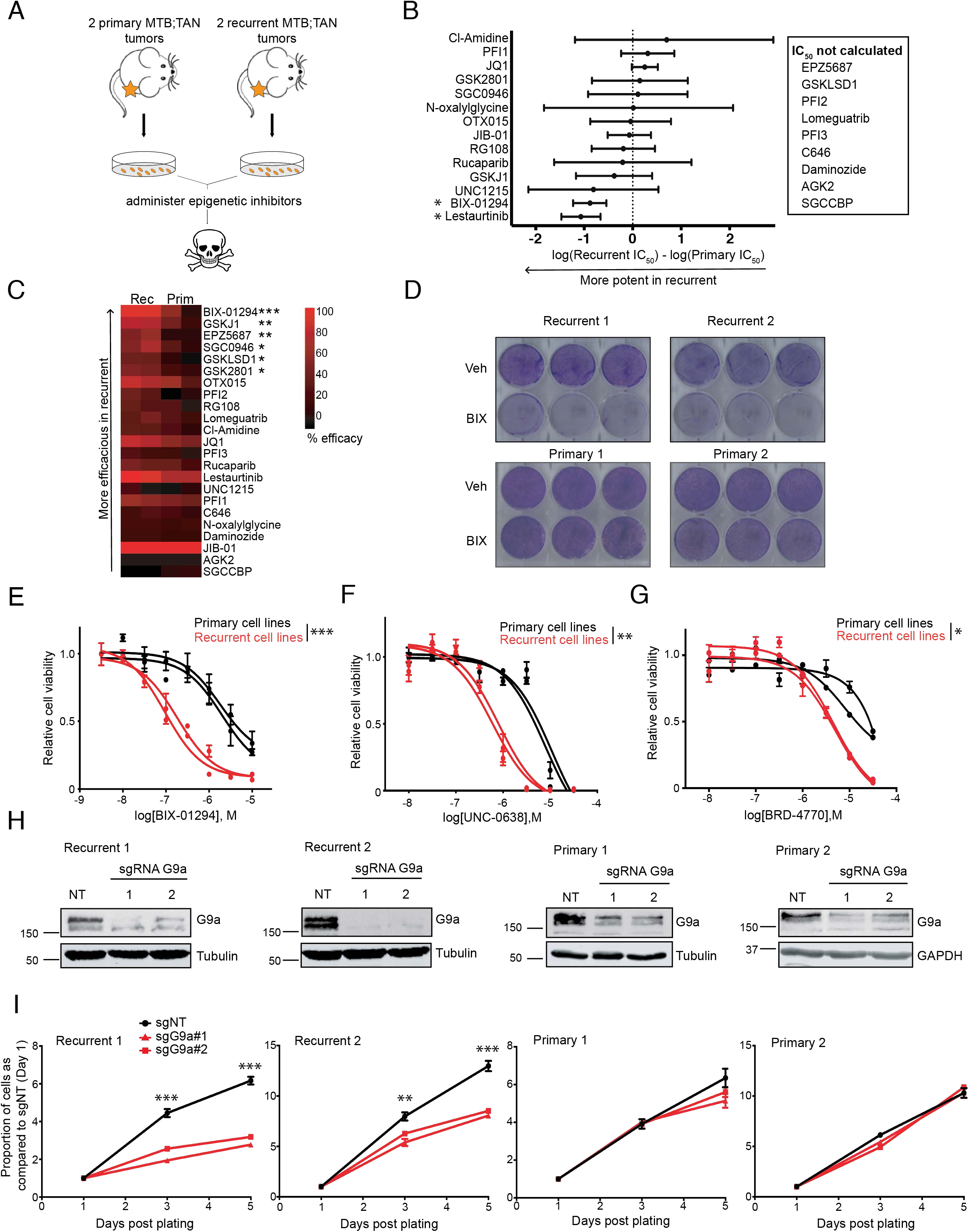
Recurrent tumors are dependent upon G9a histone methyltransferase activity. A) Tumor cells derived from primary or recurrent MTB;TAN tumors were treated with a panel of small-molecule inhibitors targeting epigenetic enzymes. B) Forest plot showing the difference in potency (i.e. IC50) between recurrent and primary tumor cells for each epigenetic inhibitor. Differences in IC50 values are shown with 95% confidence intervals. C) Heatmap showing the efficacy of each tested drug at the lowest concentration producing maximal cell growth inhibition. Significance between drug efficacy was determined by two-way ANOVA (Drug x Cohort) and Sidak’s multiple comparison test. D) Crystal violet staining of primary or recurrent tumor cells treated with vehicle or 2 µM BIX-01294 for 48 hours. E-G) Concentration response curves for primary (black) and recurrent (red) tumor cells treated with increasing concentrations of BIX-01294 (E), UNC-0638 (F), or BRD-4770 (G). IC50 and standard error values were calculated for each cohort by non-linear regression and significance was evaluated by Student’s unpaired t-test. H) Western blot analysis for G9a expression following infection with one of two independent sgRNAs targeting G9a (sgG9a#1 or sgG9a#2), or a non-targeting sgRNA (sgNT) in Cas9-expressing primary and recurrent tumor cell lines. I) Growth curves for control and G9a-knockdown primary and recurrent tumor cells. Asterisks denote significance between control and the nearest G9a sgRNA. Significance was determined by two-way ANOVA (time x sgRNA) followed by Sidak’s multiple comparison test. Error bars denote mean ± SEM. *p-value <0.05, ** p-value <0.01, *** p-value <0.001.

We focused on the G9a inhibitor BIX-01294, as it was among the most potent and efficacious inhibitors in recurrent tumor cells as compared to primary tumor cells. G9a is a histone methyltransferase that heterodimerizes with the structurally related G9a-like protein (GLP) to catalyze mono- and dimethylation of H3K9 (Jenuwein, 2006; Jenuwein et al., 1998; Tachibana et al., 2005). H3K9me1 and H3K9me2 are associated with transcriptional silencing of genes in euchromatin, while H3K9me3 – which is deposited by Setdb1 or Suv39H1 – is associated with repressed genes in heterochromatin (Rice et al., 2003; Schultz et al., 2002; Wang et al., 2003). BIX-01294 inhibits G9a/GLP by competing for binding with the amino acids N-terminal of the substrate lysine residue (Chang et al., 2009). We first confirmed that BIX-01294 selectively inhibits recurrent tumor cell growth by additional cell viability assays (Figure 2D **and** E). At high concentrations (> 4 µM), BIX-01294 also inhibits growth of primary tumor cells, possibly due to off-target effects (Kubicek et al., 2007). Therefore, we tested whether recurrent tumor cells are differentially sensitive to additional G9a inhibitors. Both UNC0638, a substrate-competitive inhibitor (Figure 2F), and BRD4770, an S-adenosylmethionine (SAM)-competitive inhibitor (Figure 2G), selectively inhibited the growth of recurrent tumor cells.

We next asked whether BIX-01294 inhibits H3K9 methylation equivalently between primary and recurrent tumor cells. BIX-01294 led to a 68-86% reduction in H3K9me2 levels in primary and recurrent tumor cells **(Figure S2A**), similar to the reduction in bulk H3K9me2 levels that has previously been reported with BIX-01294 (Vedadi et al., 2011). Furthermore, H3K9me2 reductions are seen particularly at the 300 nM and 1 µM concentrations **(Figure S2B and C)**, which is in line with the IC_50_ values observed in recurrent tumor cells.

To assess whether the effect of G9a inhibitors on recurrent cell viability was mediated through on-target inhibition of G9a, we used CRISPR-Cas9 to knock out G9a in primary and recurrent tumor cell lines. Cells were infected with lentivirus expressing a control sgRNA targeting the Rosa26 locus (sgNT), or one of two independent sgRNAs targeting G9a (sgG9a) (Figure 2H). G9a knockout significantly inhibited the growth of recurrent tumor cell lines, while primary tumor cells lines were unaffected (Figure 2I), providing genetic confirmation that recurrent tumor cells are dependent on G9a activity.

Finally, we asked whether Met-amplified recurrent tumors were also sensitive to G9a inhibitors. To address this, we tested the effect of BIX treatment on an expanded panel of 3 primary tumor cell lines and 5 recurrent tumor cell lines. While the growth of all non-Met amplified recurrent cell lines was inhibited by BIX, neither of the two Met-amplified cell lines (#4 and #5) was sensitive to BIX **(Figures S2D and E)**, indicating that Met-amplified tumors are not dependent upon G9a activity. Taken together, these results suggest that a subset of recurrent tumors acquire a dependence upon G9a methyltransferase activity.

### G9a Promotes Tumor Recurrence in Vivo

Having established that recurrent tumors are dependent upon G9a activity, we next asked whether G9a expression is altered in recurrent tumors. We measured G9a expression in an independent cohort of 7 primary and 7 recurrent MTB;TAN tumors. G9a has a long isoform (G9a long) and a short isoform (G9a short) generated by alternative splicing. There was a trend toward increase expression of the short isoform recurrent tumors (Figure 3A), and in recurrent tumor cell lines (Figure 3B). Intriguingly, mRNA levels of G9a were not increased in recurrent tumors or recurrent tumor cells **(Figures S2F and G)**, suggesting that increased G9a protein expression is not mediated through increased transcript levels.

**Figure 3:**
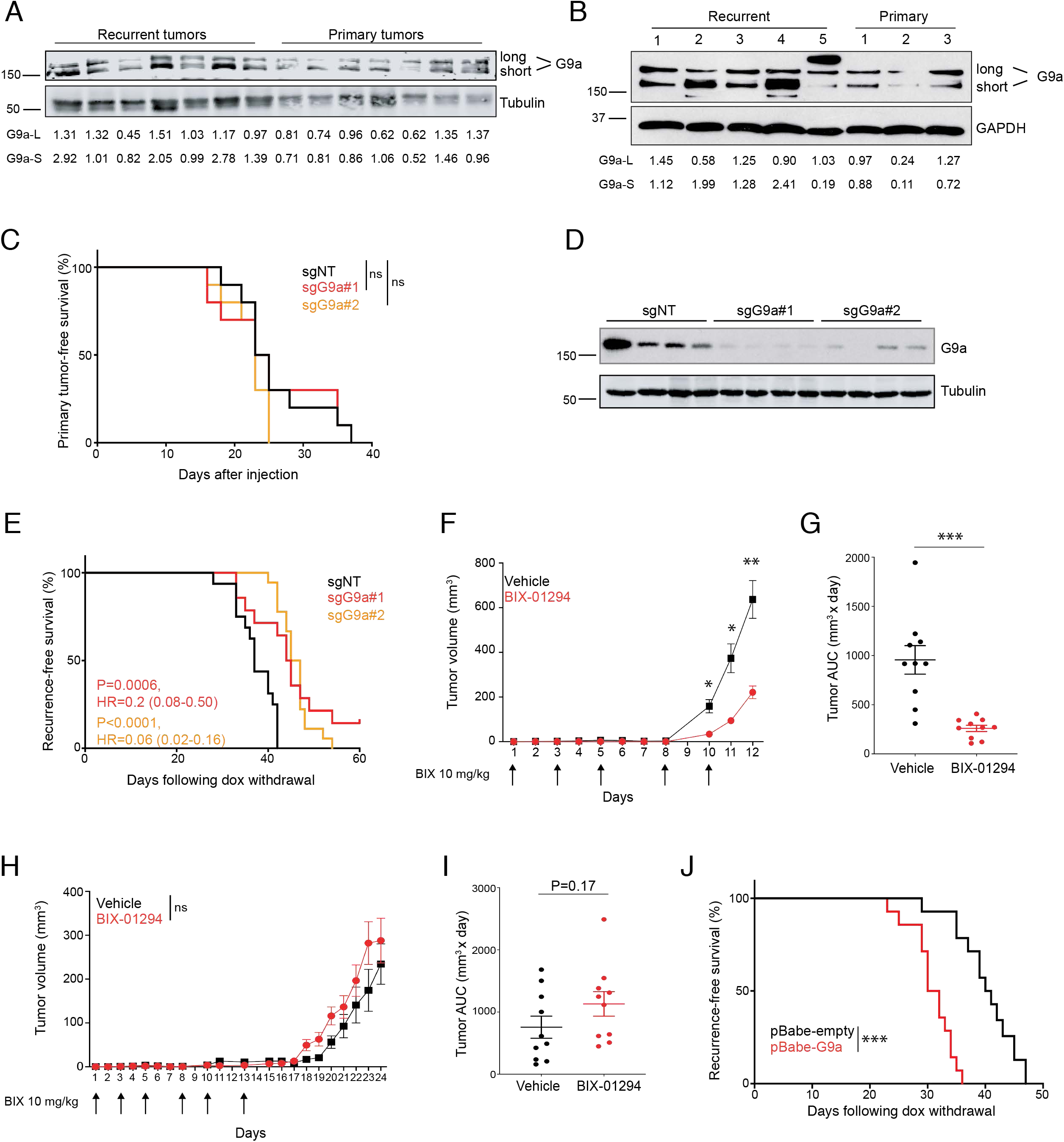
G9a promotes tumor recurrence in vivo. A) Western blot analysis for G9a expression in primary (n=7) and recurrent (n=7) MTB;TAN tumors. Quantification of each G9a isoform is shown relative to primary tumor #1. B) Western blot analysis for G9a expression in primary (n=3) and recurrent (n=5) tumor-derived cell lines. Quantification of each G9a isoform is shown relative to primary cell line #1 is shown. C) Kaplan-Meier survival curves showing time until primary tumor formation (∼75 mm^3^) for mice injected orthotopically with sgNT, sgG9a#1, or sgG9a#2-expressing primary tumor cells (primary #2; n=10 mice / cohort). Statistical significance was determined by Mantel-Cox log-rank test. D) Western blot analysis of G9a expression in representative primary tumors from panel (C). E) Kaplan-Meier survival curves showing recurrence-free survival for sgNT, sgG9a#1, or sgG9a#2-expressing tumors. P values, hazards ratios, and 95% confidence intervals are indicated as compared to sgNT. Statistical significance was determined by Mantel-Cox log-rank test. F) Mean tumor growth curves for recurrent tumor cell line #3 injected bilaterally into the mammary gland of FVB mice (n=10 tumors / cohort) and treated with vehicle or 10 mg/kg BIX-01294 three times a week. Arrows indicate drug treatments. Significance was determined by repeated-measures 2-way ANOVA (Time x Treatment) with Sidak’s post-hoc test. G) Area under the curve (AUC) values for tumor growth curves shown in (F). Significance was determined by Student’s unpaired t-test. H) Mean tumor growth curves for primary tumor cell line #2 injected bilaterally into the mammary gland of TAN mice (n=10 tumors / cohort) and treated with vehicle or 10 mg/kg BIX-01294 three times a week for two weeks. Arrows indicate drug treatments. Significance was determined by repeated-measures 2-way ANOVA (Time x Treatment) with Sidak’s post-hoc test. I) Area under the curve (AUC) values for tumor growth curves shown in (H). Significance was determined by Student’s unpaired t-test. J) Kaplan-Meier survival curves showing recurrence-free survival for control or G9a-overexpressing tumors. Statistical significance was determined by Mantel-Cox log-rank test. Results from A are representative of three independent experiments. Results from B are representative of two independent experiments. Error bars denote mean ± SEM. *P < 0.05, ** P<0.01, ***P<0.001.

The observation that recurrent tumors upregulate G9a and are dependent upon G9a activity suggested that G9a may promote the development of recurrent tumors. To test this directly, we used an orthotopic recurrence assay to determine whether G9a knockout in primary tumors affects the latency of tumor recurrence. Control or G9a-knockout cells were injected into the inguinal mammary gland of nude mice on dox to generate primary tumors. Primary tumors from control and G9a knockout cells formed with similar kinetics (Figure 3C **and S2H)** and primary tumors maintained G9a knockout (Figure 3D), consistent with our findings that G9a knockout does not affect primary tumor cell growth *in vitro*. Once primary tumors reached ∼75mm^3^, dox was withdrawn to induce Her2 withdrawal and tumor regression, and mice were monitored for the formation of recurrent tumors. G9a knockout significantly delayed the time to recurrence (*P*=0.0006, HR=0.02 (CI: 0.08-0.50) for sgG9a#1; *P*<0.0001, HR=0.06 (CI: 0.02-0.16) for sgG9a#2) (Figure 3E). Interestingly, many recurrent tumors maintained G9a knockout **(Figure S2I)**, suggesting that these tumors had bypassed the requirement for G9a activity. Given that Met-amplified are not sensitive to G9a inhibition, we considered Met amplification as one potential bypass mechanism. Indeed, we found that a greater proportion of G9a knockout recurrent tumors had amplified Met as compared to control recurrent tumors **(Figure S2J)**, though this did not reach statistical significance, likely due to the small sample size. Taken together, these data show that G9a promotes tumor recurrence following Her2 downregulation.

Next, we evaluated whether pharmacologic inhibition of G9a can inhibit the growth of existing tumors *in vivo*. Recurrent or primary tumor cells were injected orthotopically into the inguinal mammary glands of FVB mice to generate orthotopic tumors, and mice were treated with BIX-01294 (10 mg/kg, IP) three times weekly for two weeks. Consistent with *in vitro* data, BIX administration significantly reduced tumor growth and tumor burden in recurrent tumor cells (Figures 3F **and** G). In contrast, primary tumor growth was not inhibited by BIX-01294 treatment, and in fact BIX-treated primary tumors grew slightly faster than control tumors (Figures 3H **and** I). BIX treatment also slowed the growth of orthotopic recurrent tumors in athymic nude recipients **(Figure S2K)**, suggesting that the effects of G9a inhibition do not require an adaptive immune system.

Finally, we asked whether overexpression of G9a in primary tumors could accelerate tumor recurrence. Primary tumor cells were transduced with a retrovirus expressing G9a or empty vector **(Figure S2L)** and orthotopically injected into mice. Control and G9a-expressing primary tumors formed with similar kinetics (data not shown). Dox was withdrawn to induce Her2 downregulation and tumor regression, and mice were monitored for the formation of recurrent tumors. G9a expression significantly accelerated the formation of recurrent tumors (Figure 3J), further indicating that G9a promotes tumor recurrence following Her2 downregulation.

### Integrated Epigenetic and Transcriptional Analysis of G9a-regulated Genes in Recurrent Tumors

We next explored the mechanistic basis for the dependence of recurrent tumors on G9a activity by identifying genes whose expression is directly regulated by G9a specifically in recurrent tumor cells. To do this, we first performed RNA-seq on primary and recurrent tumor cells treated with BIX-01294 for 16 hours. Importantly, this early time-point precedes the induction of cell death in response to BIX-01294 treatment. Because Met-amplified recurrent tumors were not sensitive to G9a inhibition, we focused on gene expression changes induced by G9a inhibition in non-Met amplified recurrent tumor cells.

G9a inhibition led to only modest changes in gene expression in primary tumor cells (Figure 4A **and Table S2).** In contrast, G9a inhibition induced widespread changes in gene expression in recurrent tumor cells (Figure 4A). We examined genes that were more significantly altered by BIX treatment in recurrent tumor cells as compared to primary tumor cells. 306 genes were differentially upregulated following BIX treatment in recurrent tumor cells, and 137 genes were differentially downregulated following BIX treatment (Figure 4A). Among the differentially upregulated genes were genes known to induce cell cycle arrest and cell death, including p21, Gadd45a, and Ccng2 (Figure 4B **and Figure S3A)**. Importantly, the majority of genes whose expression changed following G9a inhibition were upregulated, consistent with G9a’s role in repressing genes through H3K9 methylation.

**Figure 4:**
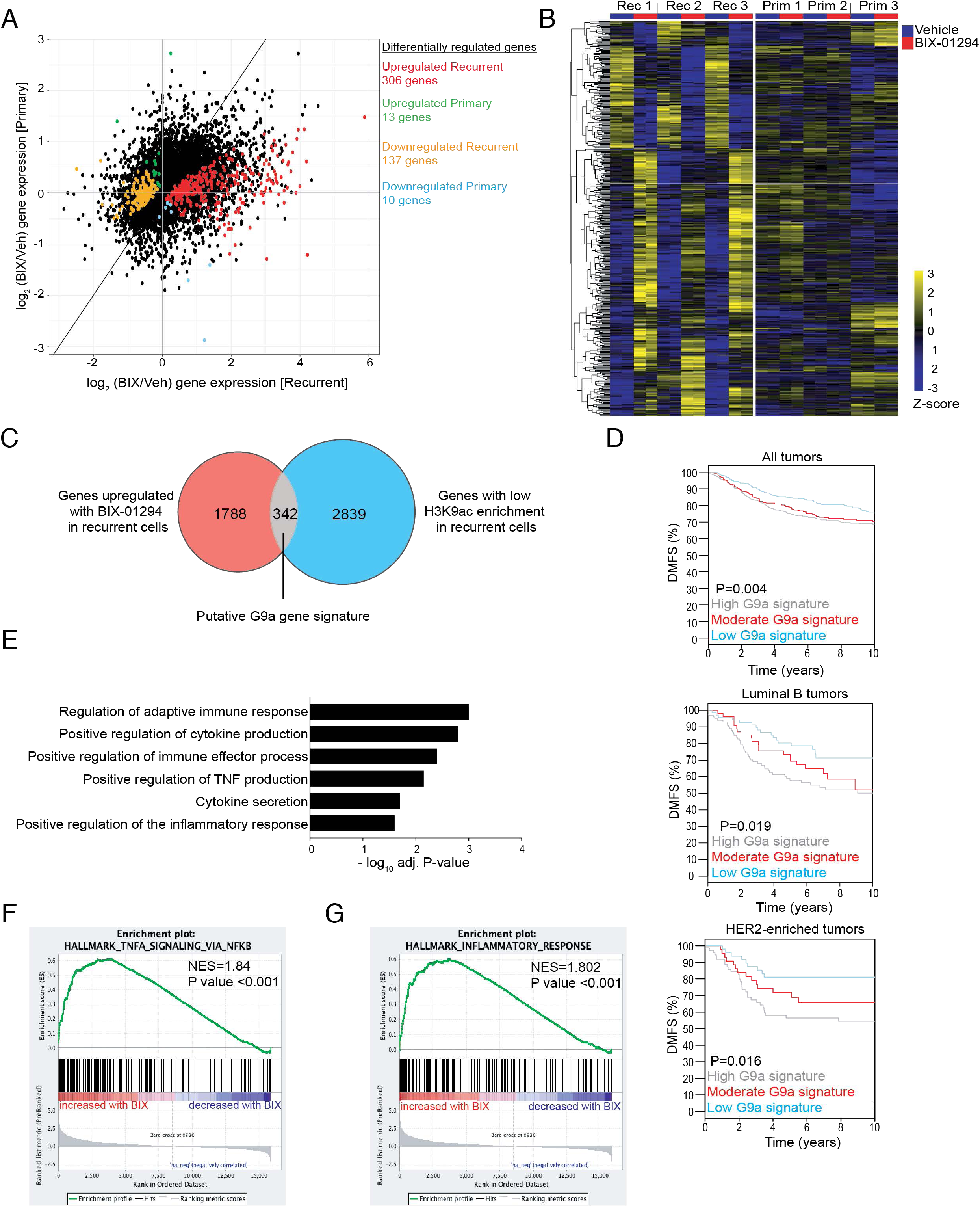
Integrated epigenetic and transcriptional analysis of G9a-regulated genes in recurrent tumors. A) Comparison of gene expression changes 16 hours after BIX-01294 (1 µM) treatment in recurrent (x-axis) and primary (y-axis) tumor cells. Colored dots indicate genes whose expression was differentially regulated by BIX treatment in primary vs. recurrent tumor cells (adjusted P-value <0.05). B) Heatmap showing median-centered, Z-score normalized expression changes for differentially genes from (A). C) A G9a gene signature was generated by overlapping genes upregulated following BIX-01294 treatment in recurrent tumor cells (adjusted P-value <0.05) with genes whose promoters had significantly lower H3K9ac in recurrent tumor cells (adjusted P-value <0.05). D) Kaplan-Meier plots showing distant metastasis-free survival (DMFS) for all tumors (n=1379), HER2-enriched tumors (n=105), and Luminal B tumors (n=225) in patients stratified by high (gray), moderate (red), or low (blue) expression of G9a signature genes. E) Gene ontology analysis showing pathways enriched in the 342 gene G9a signature from (C). F-G) GSEA plots showing enrichment of a TNF/NFκB signature (F) and an inflammatory signature (G) in recurrent tumor cells following G9a inhibition.

We next overlapped genes upregulated following G9a inhibition in recurrent tumor cells with H3K9 ChIP-seq data to generate a list of genes likely to be directly regulated by G9a through H3K9 methylation. G9a can regulate gene expression by depositing one or two methyl groups on H3K9 (Jenuwein, 2006; Jenuwein et al., 1998), and H3K9me2 can subsequently be converted to H3K9me3 by other SET domain-containing methyltransferases, including Setdb1 and Suv39H1 (Rice et al., 2003; Schultz et al., 2002; Wang et al., 2003). Because we were unable to successfully perform ChIP-sequencing to identify H3K9me2 peaks (data not shown), we considered a reduction in H3K9 acetylation in recurrent tumor cells as a proxy for increased H3K9 methylation. We identified 342 genes that (i) had lower H3K9 acetylation in recurrent tumor cells as compared to primary tumor cells, and (ii) were upregulated following G9a inhibition specifically in recurrent tumor cells (Figure 4C and **Table S3)**. We used ChIP-qPCR to assess H3K9me2 levels at the promoter of 10 of these putative G9a target genes, and found that 9/10 genes had elevated levels of H3K9me2 in recurrent tumor cells, suggesting that lack of H3K9ac is a suitable surrogate for H3K9 methylation **(Figure S3B)**. We hypothesized that these 342 genes represent putative direct targets of G9a in recurrent tumor cells whose re-expression may mediate cell death following G9a inhibition. Interestingly, this G9a-regulated gene set is correlated with increased risk of distant recurrence in human cancers, particularly among HER2-enriched and Luminal B breast tumors (Figure 4D) (Ringner et al., 2011).

Gene ontology analysis revealed that these G9a targets are enriched for genes involved in regulation of inflammatory responses, cytokine production, and tumor necrosis factor (TNF) signaling (Figure 4E). Examination of specific genes revealed that the pro-inflammatory cytokines TNF, IL-23a, and Cxcl2 were all induced following G9a inhibition in recurrent tumor cells **(Figure S3C-E)**. Consistent with this, Gene Set Enrichment Analysis (GSEA) showed enrichment of a TNF-NFκB signature and an inflammatory response signature in recurrent cells treated with BIX-01294 (Figures 4F **and** G). Taken together, these results indicate that G9a directly represses an inflammatory gene expression program in recurrent tumor cells, and raise the possibility that upregulation of pro-inflammatory genes may contribute to cell death in recurrent tumor cells following G9a inhibition.

Given that G9a inhibition induces expression of pro-inflammatory genes in recurrent tumor cells, we examined whether G9a inhibition is associated with alterations in the tumor immune microenvironment. To evaluate this, we performed immune cell profiling on recurrent and primary orthotopic tumors treated with BIX-01294 (see Figures 3F and H) using markers for leukocytes, monocytes, macrophages, and T-cells. G9a inhibition did not significantly alter any of the immune cell populations in recurrent **(Figure S3F)** or primary tumors **(data not shown)**. While these results do not rule out the possibility that G9a inhibition affects the tumor immune microenvironment, they do suggest that G9a inhibition slows the growth of recurrent tumors through tumor cell-intrinsic pathways. This is consistent with the observation that BIX treatment inhibits recurrent tumor cell growth in vitro (see Figure 2) and in immunocompromised mice (see Figure S2K and L).

### G9a-dependent Silencing of TNF is Required for Recurrent Tumor Cell Survival

To investigate whether upregulation of pro-inflammatory cytokines contributes to cell death following G9a inhibition, we focused our attention on TNF. TNF was among the most differentially induced cytokines following BIX-01294 treatment in recurrent versus primary tumor cells, and the TNF promoter had reduced H3K9 acetylation in recurrent as compared to primary tumor cells (Figure 5A). Further, G9a has been reported to regulate TNF in other models (Li et al., 2018). TNF is a pleiotropic cytokine that has both pro- and anti-tumor effects, depending on the cellular context (Rokhlin et al., 2000). TNF can induce tumor cell-specific cell death in certain cancer cell types and lineages, including breast cancer (Burow et al., 1998; Wang and Lin, 2008). We first tested whether G9a regulates TNF through H3K9 methylation in our model. At baseline, TNF expression was slightly lower in recurrent tumor cells as compared to primary tumor cells **(Figure S3A)**. G9a inhibition led to a 4- to 50-fold increase in TNF levels in recurrent cells, but only induced modest changes in TNF levels in primary cells, as measured by qRT-PCR (Figure 5B). Further, H3K9me2 levels at the TNF promoter were significantly higher in recurrent tumor cells as compared to primary tumor cells (Figure 5C), and were reduced following BIX-01294 treatment (Figure 5D), confirming that G9a promotes H3K9 methylation at the TNF promoter. We next asked whether TNF selectively inhibits recurrent tumor cell growth. Treatment with recombinant TNF led to a marked decrease in the growth of recurrent tumor cells, but had only a modest effect on primary tumor cells (Figure 5E). Similar results were obtained in long-term colony formation assays (Figure 5F). Taken together, these results demonstrate that G9a directly silences pro-inflammatory cytokines, including TNF, in recurrent tumor cells, and that re-activation of these genes mediates the anti-tumor effects of G9a inhibition in recurrent tumors.

**Figure 5:**
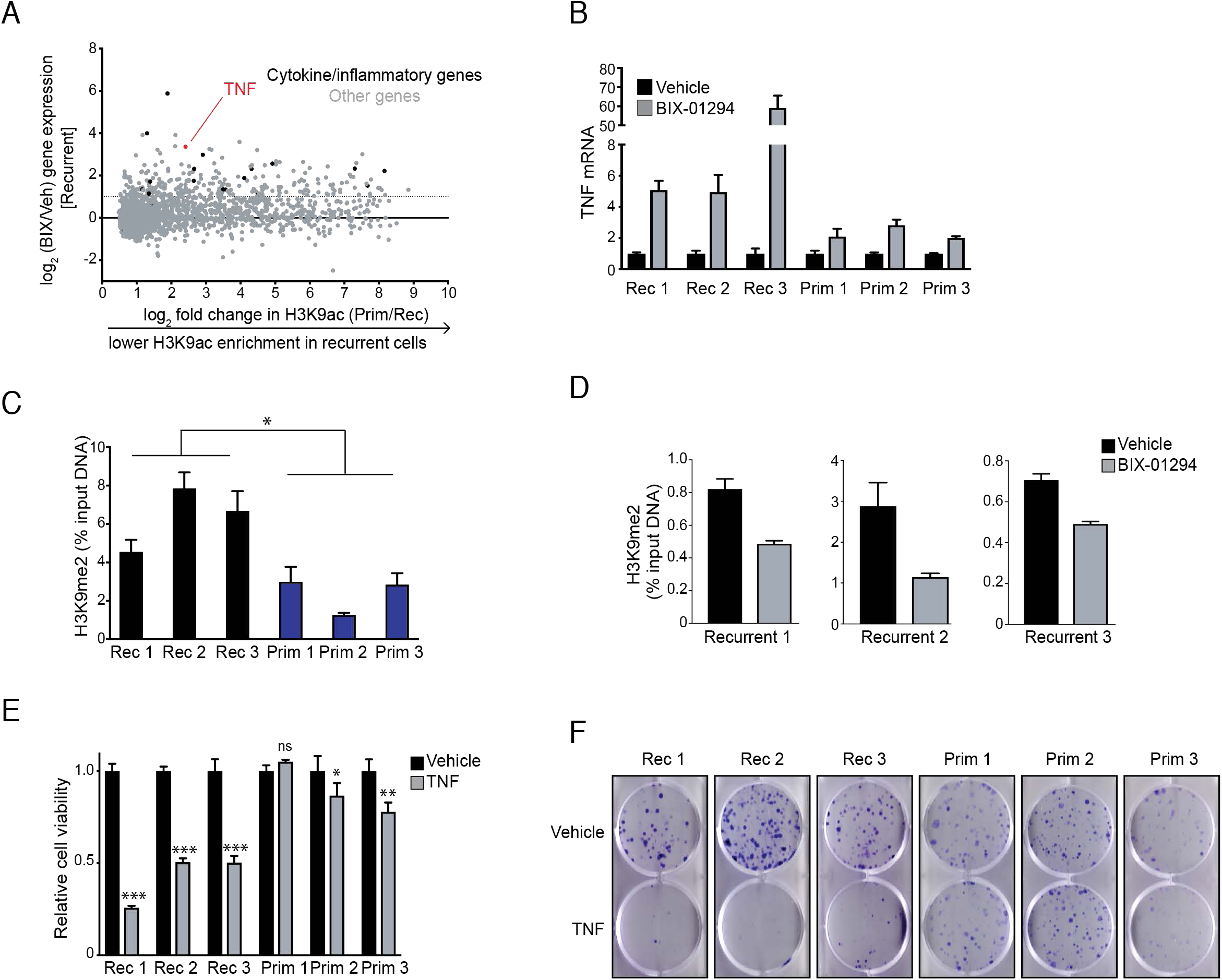
G9a-dependent silencing of TNF is required for recurrent tumor cell survival. A) Scatterplot showing genes induced following BIX-01294 treatment in recurrent tumor cells (y-axis) as a function of differential H3K9ac peaks in recurrent tumor cells (x-axis). Dashed line indicates 2-fold mRNA upregulation. Inflammatory genes identified from the G9a-regulated gene set are indicated in black. B) qPCR analysis for TNF expression 16 hours after BIX-01294 treatment (1 µM) in recurrent and primary tumor cell lines. Expression values were normalized to the vehicle within each cell line. C) ChIP-qPCR showing H3K9me2 enrichment at the TNF promoter in recurrent and primary tumor cells. D) ChIP-qPCR showing H3K9me2 enrichment at the TNF promoter in recurrent cell lines following 16 hours of 1 µM BIX-01294 treatment. E) Cell viability of primary and recurrent tumor cells treated with TNF (10 ng/mL) for 3 days. Significance was determined by one-way ANOVA and Tukey’s post-hoc test. F) Representative colony formation assays showing viability of primary and recurrent tumor cells after 7-day treatment with TNF (10 ng/mL). Results from panels E-F are representative of at least two independent experiments. Error bars denote mean ± SEM. *p<0.05, **p<0.001***p<0.0001, ns=not significant.

### G9a Inhibition Leads to Induction of p53 Targets and p53-dependent Cell Death

RNA-seq analysis revealed that a p53 signature was enriched following G9a inhibition **(Figure S4A)**. It has been previously described that BIX-01294 mediates cell death in human cancer cell lines through a p53-dependent mechanism (Fan et al., 2015). Furthermore, p53 plays a crucial role in TNF-mediated cell death (Pastor et al., 2010). Given these reports, we hypothesized that p53 may be required for cell death following G9a inhibition in recurrent tumor cells. We first confirmed that p53 was expressed and not mutated in recurrent tumor cells. Western blot analysis showed that p53 protein is expressed in all primary and recurrent tumor cell lines **(Figure S4B)**. Also, sequencing of p53 identified no mutations in either primary or recurrent tumor cell lines (data not shown). We next confirmed that p53 targets were upregulated following treatment with BIX-01294. qPCR analysis showed that the canonical p53 targets Cdkn1a (p21) and Gadd45a were significantly upregulated in BIX-treated recurrent tumor cell lines **(Figure S4C).** Together, these data suggest that the p53 signaling axis is intact in recurrent tumor cells and that p53 targets genes are upregulated in response to G9a inhibitors.

To determine whether p53 is required to mediate cell death in response to G9a inhibition, we knocked out p53 in recurrent tumor cell lines using CRISPR/Cas9 **(Figure S4D)**. p53 knockout markedly reduced the induction of p21 and Gadd45a following BIX treatment in recurrent tumor cell lines **(Figure S4E)**. Surprisingly, the induction of p21 and Gadd45a was not completely reduced following p53 knockdown, suggesting that G9a may regulate expression of these genes in part through p53-independent mechanisms, potentially through methylation of H3K9. Significantly, p53 knockout resulted in an approximately 3-fold shift in the potency of BIX-01294 in recurrent tumor cell line #1 (IC_50_ for sgNT 168 nM [95% CI: 102 nM – 273 nM]; IC_50_ for sgp53 613 nM [95% CI: 345 nM – 996 nM]) and an approximately 2-fold shift in recurrent cell lines #2 (IC_50_ for sgNT 120 nM [95% CI: 79 nM – 182 nM]; IC_50_ for sgp53 248 nM [95% CI: 159 nM – 386 nM]) and #3 (IC_50_ for sgNT 476 nM [95% CI: 304 nM – 745 nM]; IC_50_ for sgp53 867 nM [95% CI: 535 nM –1.4 µM]) **(Figure S4F)**. Consistent with this, Annexin V staining showed that p53 knockout partially decreased cell death following BIX-01294 treatment **(Figure S4G)**. Taken together, these results indicate that p53 partially contributes to cell death following G9a inhibition, suggesting the presence of both p53-dependent and -independent cell death pathways.

### G9a Inhibition Induces Necroptotic Cell Death

The results described above indicate that G9a inhibition leads to upregulation of pro-inflammatory cytokines, and one such cytokine, TNF, is sufficient to reduce cell viability in recurrent cells. Because TNF can induce both apoptotic and necroptotic cell death pathways (Laster et al., 1988; Nikoletopoulou et al., 2013), we next sought to define the mode of cell death following G9a inhibition. G9a inhibition led to a marked increase in Annexin V staining only in recurrent tumor cells (Figure 6A-B). Annexin V binds to phosphatidylserine (PS) on the outer leaflet of the plasma membrane. PS flipping is a marker of both apoptosis and necroptosis, and so to differentiate between these cell death pathways we examined molecular markers specific for each cell death pathway. Surprisingly, the apoptotic markers cleaved caspase-3 and cleaved PARP were not induced following BIX-01294 treatment (Figure 6C). Instead, BIX-01294 treatment led to a marked increase in phosphorylation of S345 of MLKL (Figure 6C), which is a marker of necroptosis and is associated with induction of necroptosis during TNF-mediated cell death (Rodriguez et al., 2016). Consistent with this, we found that a necroptosis gene expression signature containing 141 necroptosis-associated genes **(Table S4)** (Callow et al., 2018; Chen et al., 2018a; Hitomi et al., 2008; Zhu et al., 2018) was enriched following G9a inhibition in recurrent tumor cells (Figure 6D). These results indicate that G9a inhibition leads to necroptotic cell death in recurrent tumor cells.

**Figure 6:**
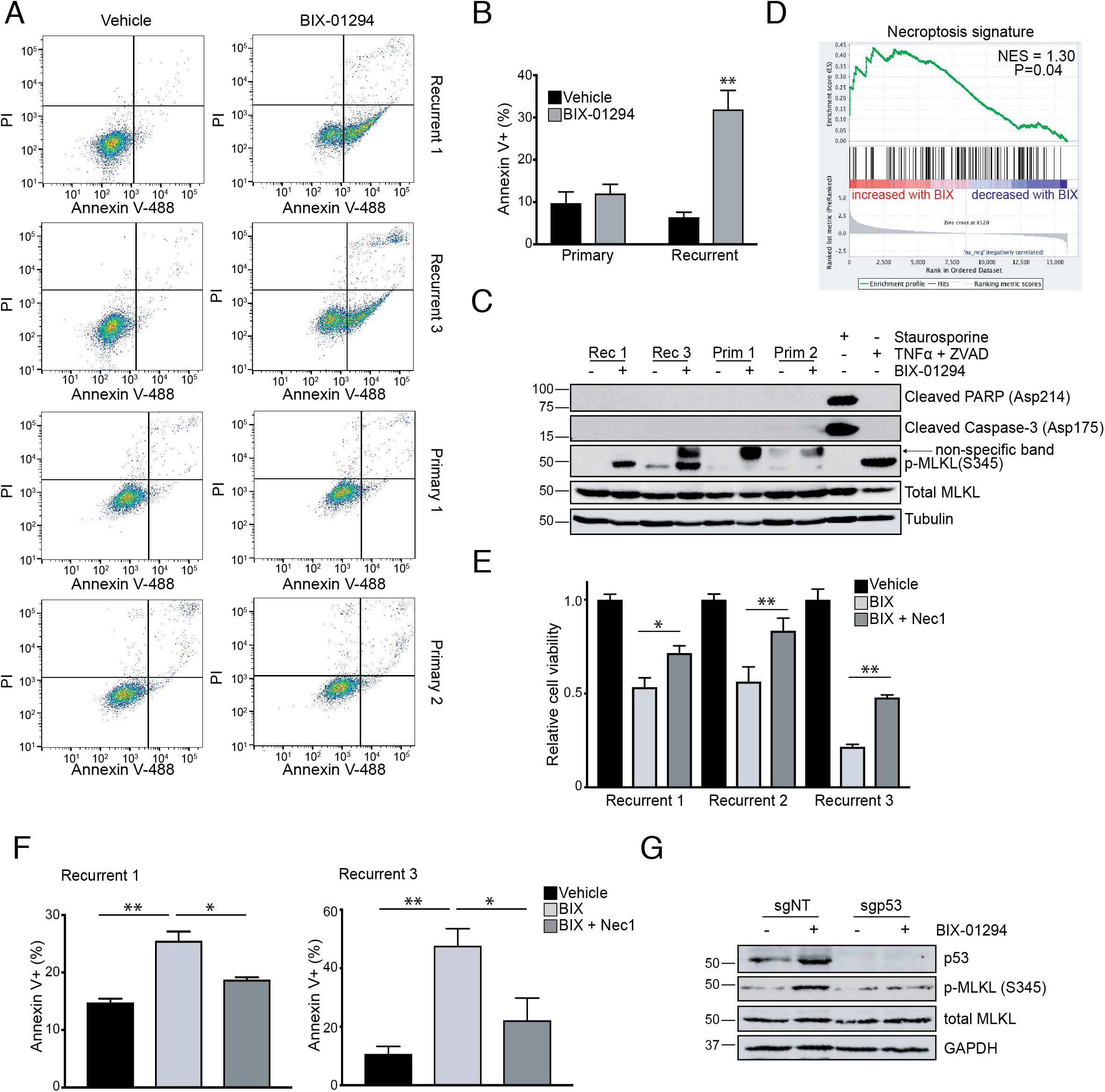
G9a inhibition induces necroptotic cell death in recurrent tumors. A) Annexin V/PI staining of recurrent and primary tumor cells treated with vehicle or 1 µM BIX-01294 for 16 hours. B) Quantification of Annexin V-positive cells from (A). Significance was determined by one-way ANOVA and Tukey’s post-hoc test. C) Western blot analysis for cleaved PARP (Asp214), cleaved Caspase-3 (Asp175), p-MLKL (S345), and total MLKL in recurrent and primary tumor cells treated with vehicle or 1 µM BIX-01294. Staurosporine and TNF + Z-VAD-FMK were included as controls for apoptosis and necroptosis, respectively. D) GSEA showing enrichment of a curated necroptosis signature in recurrent tumor cells treated with BIX-01294. P-value and normalized enrichment score are shown. E) Cell viability of recurrent tumor cells after 16-hour treatment with BIX-01294 (300 nM) alone or in combination with necrostatin-1 (30 µM). F) Quantification of Annexin V staining in recurrent tumor cells (#1) treated for 24 hours with BIX-01294 (750 nM) alone or in combination with necrostatin-1 (30 µM). Significance in (E) and (F) was determined by one-way ANOVA with Tukey’s posthoc test. G) Western blot analysis showing p53, p-MLKL (S345), and total MLKL in control or p53-knockout recurrent tumor cells (#3) treated with vehicle or16 hours with 1 µM BIX-01294. Results from A-C, E-G are representative of at least two independent experiments. Error bars denote mean ± SEM. *p<0.05, **p<0.01, ns=not significant.

To evaluate whether induction of necroptosis is necessary for BIX-mediated cell death, we pre-treated recurrent tumor cell lines with the RIPK1 inhibitor Necrostatin-1 (Nec-1). Nec-1 partially rescued cell viability following BIX treatment in three recurrent tumor cell lines (Figure 6E). Consistent with this rescue of cell viability, Nec-1 also decreased the proportion of Annexin V-positive cells following BIX-01294 (Figure 6F). Taken together, these results indicate that G9a inhibition induces necroptotic-dependent cell death in recurrent tumor cells.

In light of our previous observation that p53 contributes to cell death following G9a inhibition, we next addressed the role of p53 in BIX-induced necroptosis. p53 has been suggested to contribute to necroptosis through indirect upregulation of the critical necroptosis intermediates RIPK1, RIPK3 and cathepsins (Tu et al., 2009; Wang et al., 2016). To address this possibility, we compared the induction of necroptosis following G9a inhibition in control and p53 knockout cells. p53 knockout cells failed to induce MLKL phosphorylation following BIX-01294 treatment (Figure 6G). Consistent with this, qPCR analysis indicated that TNF upregulation was blunted in p53 knockout cells treated with BIX, suggesting that p53 activity is required for TNF expression **(Figure S4H)**. Taken together, our data suggest that p53 is required for TNF-dependent necroptosis in recurrent tumor cells.

### Recurrent Tumor Cells are Dependent on RIPK3

Our findings to this point demonstrated that G9a inhibition induces inflammatory cytokine expression, leading to p53-dependent necroptosis. Paradoxically, primary tumor cell lines demonstrated similar p53 expression as recurrent tumor cell lines, but tolerated exogenous TNF. Therefore, we postulated that recurrent tumor cells acquire an enhanced sensitivity to necroptosis in response to exogenous stimuli that is lacking in primary tumor cells.

To address this question, we evaluated the expression of critical intermediates in the necroptosis pathway. Necroptosis is triggered through a multiprotein complex called the necrosome consisting of RIPK1 and RIPK3 (Newton, 2015), which acts to phosphorylate and activate MLKL, ultimately leading to cell membrane disruption and necroptosis (Weinlich et al., 2017). Conversely, caspase-8 can inhibit necroptosis at least in part through cleaving RIPK1 and RIPK3 (O’Donnell et al., 2011). To understand why recurrent tumors have heightened sensitivity to necroptosis, we examined expression of RIPK1, RIPK3, caspase-8 and MLKL in primary and recurrent tumor cells (Figure 7A **and Figures S5A-C)**. RNA sequencing revealed that RIPK1 and MLKL were expressed at similar levels between primary and recurrent tumor cells **(Figures S5B and C)**. In contrast, we observed a 2-fold decrease in caspase-8 expression and a nearly 1000-fold increase in RIPK3 expression in recurrent tumor cells (Figure 7A **and Figure S5A)**. qPCR and Western blotting confirmed that RIPK3 was dramatically upregulated in recurrent tumor cells relative to primary tumor cells (Figure 7B **and** C). Intriguingly, Met-amplified recurrent tumor cell lines did not upregulate RIPK3, consistent with the finding that these cells are resistant to G9a inhibition (Figure 7B **and** C). To test whether RIPK3 is sufficient to induce sensitivity to TNF-induced necroptosis, we engineered primary tumor cells expressing RIPK3. RIPK3 expression in primary tumor cells increased their sensitivity to TNF-induced cell death **(Figure S5D-E)**. These results suggest that upregulation of RIPK3 in recurrent tumor cells underlies, at least in part, their sensitivity to both G9a inhibition and TNF treatment.

**Figure 7:**
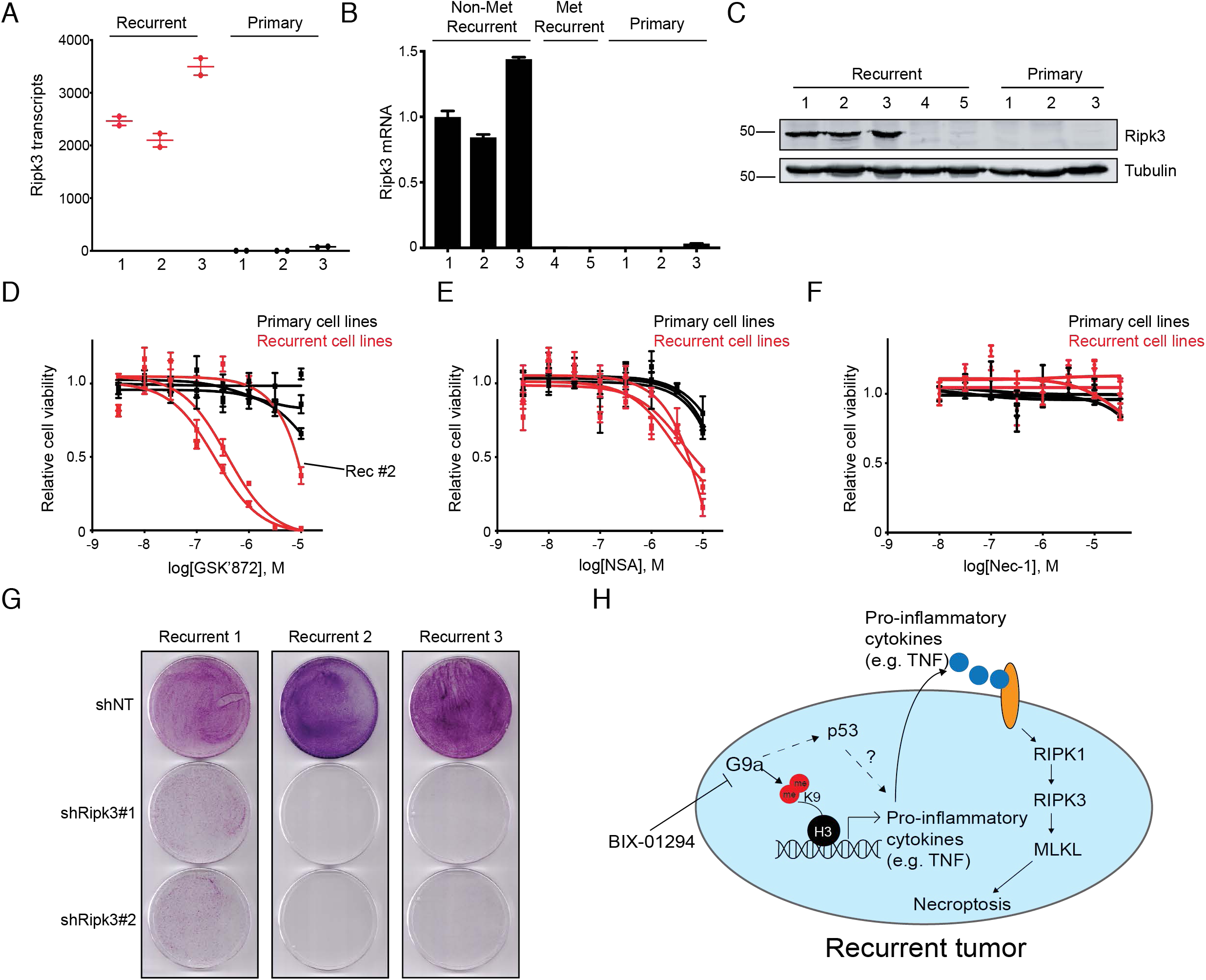
Recurrent tumors are dependent on RIPK3. A) RNA transcript counts for RIPK3 in recurrent and primary tumor cells as determined by RNA-seq analysis. B) qPCR showing RIPK3 expression in non-Met amplified recurrent tumor cells, Met-amplified recurrent tumor cells, and primary tumor cells. C) Western blot analysis for RIPK3 expression in primary and recurrent tumor cells. D-F) Concentration-response curves for primary and recurrent tumor cells treated with increasing doses of GSK’872 (D), Necrosulfonamide (E), and Necrostatin-1 (F). G) Cell viability following genetic knockdown of RIPK3 with two independent shRNAs or a scrambled control shRNA. H) RIPK3-driven recurrent tumors require G9a activity to silence inflammatory cytokine expression and prevent necroptosis. Error bars denote mean ± SEM.

RIPK3 is traditionally thought to function as a tumor suppressor (Vucur et al., 2013). Consistent with this, RIPK3 expression is silenced in >85% of breast cancer patients (Koo et al., 2015). In contrast to its tumor suppressive function, several recent reports have suggested that RIPK3 may drive tumor growth (Hanggi et al., 2017; Liu et al., 2016; Seifert et al., 2016). Given that RIPK3 is highly expressed in recurrent and not primary tumor cells, we hypothesized that recurrent tumor cells may depend on RIPK3 activity. To test this hypothesis, we tested the effect of pharmacologic inhibition of RIPK3 on tumor cell growth; inhibitors against RIPK1 and MLKL were included as controls. Whereas the RIPK1 inhibitor Nec-1 and the MLKL inhibitor NSA had minimal effects on recurrent tumor cell viability, the RIPK3 inhibitor GSK’872 profoundly inhibited recurrent tumor cell viability in two cell lines (Figure 7D-F). Further, all three recurrent cell lines exhibited decreased cell viability in response to genetic knockdown of RIPK3 with two independent shRNAs (Figure 7G **and Figure S5F)**. Taken together, these data suggest that elevated RIPK3 expression promotes recurrent tumor cell growth, and that this can be pharmacologically targeted.

Finally, we investigated whether G9a also regulates cell viability and inflammatory cytokine expression in human breast cancer cell lines. To address this, we measured the response of 24 breast cancer cell lines to increasing concentrations of BIX-01294; the normal mammary epithelial cell line MCF10A was included as a control **(Figure S5G and H)**. Only SKBR3 cells had an IC_50_ for BIX below 1 µM, consistent with previous reports that SKBR3 cells are sensitive to G9a inhibition **(Figure S5G and H)**(Kim et al., 2013). Similar to recurrent mouse tumors, SKBR3 cells had elevated expression of RIPK3 as compared to MCF7 and MCF10A cell lines, both of which are resistant to G9a inhibition **(Figure S5I)**. Further, BIX treatment induced robust expression of TNF in multiple breast cancer cell lines **(Figure S5J)**. These data suggest that G9a can regulate TNF expression in a range of human breast cancer cell lines.

## Discussion

The development of targeted therapies against Her2 has been a major advance in the treatment of Her2-positive cancers. However, a fraction of Her2-positive tumors treated with adjuvant therapies will eventually recur, representing one of the largest hurdles to obtaining cures in breast cancer (Haque et al., 2012; Sparano et al., 2015). Many Her2-positive tumors become Her2-negative or refractory to anti-Her2 therapies upon relapse, suggesting that these tumors have acquired dependence upon alternative pathways for their growth (Hurley et al., 2006; Mittendorf et al., 2009). In the current study, we used a genetically engineered mouse model of Her2-driven breast cancer to examine mechanisms underlying Her2-independent tumor recurrence. We found that a subset of recurrent tumors underwent widespread epigenetic reprogramming and displayed profound changes in gene expression as compared to primary tumors. These epigenetic changes were associated with an acquired dependency of recurrent tumors on the histone methyltransferase G9a. Genetic knockout of G9a delayed tumor recurrence, and pharmacologic inhibition of G9a prevented the growth of recurrent tumors. Mechanistically, G9a was required in recurrent tumors for the silencing of pro-inflammatory genes. Inhibition of G9a led to the re-expression of pro-inflammatory genes and the induction of necroptotic cell death. Surprisingly, recurrent tumors had dramatically upregulated expression of the essential necroptosis protein RIPK3. RIPK3 activity was both required for recurrent tumor cell growth and sensitized recurrent tumor cells to necroptosis. Taken together, we found that G9a-dependent epigenetic reprogramming promotes breast cancer recurrence, and identified a G9a-RIPK3 pathway as a targetable collateral vulnerability in recurrent breast cancer (Figure 7H).

RNA-seq and ChIP-seq analysis revealed that recurrent tumors had profound transcriptional and epigenetic differences from primary tumors. Results from an epigenetic inhibitor screen suggested that recurrent tumors had acquired novel epigenetic dependencies as compared to primary tumors. Together, this strongly suggests that epigenetic remodeling is functionally important for tumor recurrence. Indeed, inhibition of G9a delayed tumor recurrence and slowed recurrent tumor growth. While the results presented here focus on G9a, it is possible that other chromatin-modifying enzymes are also functionally important for tumor recurrence. Consistent with this, we found that recurrent tumor cells had acquired novel H3K9ac, H3K4me3 and RNApol2 peaks at intergenic regions. Although the mechanisms underlying these epigenetic changes remain unknown, our results suggest that epigenetic remodeling is a major feature of tumor recurrence, and this remodeling is associated with an acquired dependence on the G9a histone methyltransferase. While our study focused on how G9a regulates gene expression through alterations in histone methylation, it is important to note that G9a has non-histone substrates, including p53 (see below), Reptin (Lee et al., 2010), and Pontin (Lee et al., 2011). G9a regulation of these pathways through direct methylation could underlie, at least in part, the dependence of recurrent tumors on G9a activity.

By integrating global gene expression and ChIP-seq data, we found that G9a directly regulates the expression of a set of pro-inflammatory genes, including TNF, in recurrent tumor cells. We found that treatment with TNF by itself was sufficient to inhibit the growth of recurrent, but not primary, tumor cells. Mechanistically, the selective sensitivity of recurrent tumor cells to TNF was due to the fact that recurrent tumor cells had upregulated the essential necroptosis kinase RIPK3. Expression of RIPK3 sensitized primary tumor cells to TNF-induced cell death, while blocking necroptosis partially reversed recurrent tumor cell death in response to G9a inhibition. RIPK3 is traditionally thought to function as a tumor suppressor, and its expression is often silenced in breast cancers (Koo et al., 2015), and so its upregulation in recurrent tumors was unexpected. However, a handful of recent reports have identified a pro-tumorigenic function for RIPK3 that is independent of necroptosis (Newton, 2015; Seifert et al., 2016). Consistent with this, we found that RIPK3 was required for the growth of recurrent tumor cells. Taken together, this suggests a model where RIPK3 upregulation is both required for recurrent tumor cell growth and sensitizes these cells to stimuli that induce necroptosis, including TNF. According to this model, the dependence of recurrent tumors for G9a activity is due, at least in part, to the requirement for recurrent tumor cells to silence TNF expression. This is an example of the concept of collateral sensitivity, where a resistance pathway – in this case high RIPK3 expression – results in enhanced sensitivity to a secondary pathway, G9a.

Our mechanistic studies suggested that the induction of necroptosis following G9a inhibition is partially dependent upon p53 activity. Other studies have shown that G9a inhibition induces p53-dependent autophagy (Fan et al., 2015), and G9a has been reported to repress p53 activity through direct methylation of lysine 373 of human p53 (K370 in mice) (Huang et al., 2010). Our data suggest that p53 acts upstream of TNF expression, thereby modulating activation of necroptosis. However, the specific role of p53 in initiating necroptosis requires further investigation. It is possible that G9a inhibition directly activates p53 by relieving an inhibitory methylation on K370. Alternatively, G9a inhibition may indirectly activate p53 through an unidentified mechanism, for instance by causing replication stress, alterations to chromatin structure, or metabolic stress (Avgustinova et al., 2018; Puzio-Kuter, 2011; Tanikawa et al., 2012). Indeed, the upregulation of p21 and Gadd45a following G9a inhibition was not completely abrogated by p53 knockout, suggesting that G9a inhibition may cause replication stress independent of p53 (Avgustinova et al., 2018). Notwithstanding the specific mechanisms, our results show that p53 activity following G9a inhibition is required for TNF upregulation and necroptosis.

A number of previous studies have found that inhibitors targeting G9a (Casciello et al., 2017; Tu et al., 2018) or the H3K27 methyltransferase Ezh2 (Chen et al., 2018b; Gardner et al., 2017) are effective in various cancers, either alone or in combination with chemotherapy. However, more recent studies have suggested that the effects of epigenetic inhibitors can be more complex, and are likely to be context-dependent. For instance, two recent studies have shown that inhibition of G9a drives the formation of tumors with more aggressive, stem-like phenotypes (Avgustinova et al., 2018; Rowbotham et al., 2018). In the study by Avgustinova et al., G9a knock out delayed the formation of carcinogen-induced skin tumors, but once G9a-knockout tumors formed they were more aggressive, with higher levels of genomic instability and more frequent p53 loss (Avgustinova et al., 2018). These results suggest that G9a can play context-dependent roles in regulating tumor progression, and underscore the importance of crosstalk between G9a and p53 in dictating the net result of G9a inhibition. Similarly, recent work in lung cancer showed that Ezh2 inhibition has anti-tumor effects in the short-term, but leads to induction of an inflammatory program – including TNF signaling – that ultimately promotes resistance to Ezh2 inhibitors (Serresi et al., 2018). The findings we present here suggest that G9a inhibition can similarly induce a pro-inflammatory program, but that this has anti-tumor rather than pro-tumor consequences. It is possible that high RIPK3 expression in recurrent tumors tilts that balance of TNF signaling toward necroptosis rather than NFκB-dependent cell survival, consistent with the dual roles of this pleiotropic cytokine (Annibaldi and Meier, 2018). Alternatively, epigenetic inhibitors may blunt tumor growth in the short-term, but lead to microenvironmental changes that ultimately drive tumor progression. Future work will be required to further define the cellular and genetic contexts in which G9a inhibitors will prove effective.

In conclusion, our results demonstrate that tumor recurrence is associated with widespread epigenetic reprogramming and acquired dependence on the histone methyltransferase G9a. These findings elucidate a novel role for G9a in silencing pro-necroptotic inflammatory cytokines in cancer. Finally, these observations suggest that strategies designed to target G9a may have clinical utility to improve survival in patients diagnosed with recurrent breast cancer.

## Materials and Methods

### Study Approval

Animal care and all animal experiments were performed with the approval of and in accordance with Duke University IACUC guidelines (A199-17-08).

### Mice

MMTV-rtTA;TetO-Her2/neu (MTB;TAN) and TetO-Her2/neu (TAN) mice are on an FVB background. Wild-type FVB mice were purchased from Jackson laboratory. Mice were fed a standard chow diet and housed under barrier conditions with 12-hour light/dark cycles.

### Tissue culture

Primary and recurrent tumor cell lines were derived and grown as previously described (Alvarez et al., 2013; Mabe et al., 2018). All human cell lines were obtained from Duke University Cell Culture Facility and cultured according to ATCC recommendations. Cell lines were authenticated and tested for mycoplasma according to standard procedures at Duke. Cell lines were used within 6 months of receipt.

The following drugs were purchased and utilized as part of the CaymanChem epigenetics screening library (#11076): Cl-Amidine, PFI-1, JQ1, GSK2801, SGC0946, N-oxalylglycine, OTX015, JIB01, RG108, Rucaparib, GSKJ1, UNC1215, lestaurtinib, BIX-01294, EPZ5687, GSKLSD1, PFI2, Lomeguatrib, PFI3, C646, Daminozide, AGK2, SGCCBP. Additional drugs included, UNC-0638 (Tocris), BIX-01294 (Tocris), BRD4770 (SelleckChem), Necrostatin-1 (SelleckChem), GSK’872 (SelleckChem), necrosulfonamide (SelleckChem), and Z-VAD-FMK (SelleckChem). Drugs were solubilized per manufacturer recommendations and utilized at concentrations stated within the text. Matching vehicle controls were used for each experiment.

### Plasmids and viral transduction

To knock out expression of G9a or p53, Cas9 was stably infected in cell lines using a lentiviral construct encoding lentiCas9-Blast (a gift from Feng Zhang, Addgene #52962). The single-guide RNAs targeting G9a and p53 listed in **Table S5** were cloned into lentiGuide-Puro (a gift from Feng Zhang, Addgene #52963). For knockdown of RIPK3, lentiviral shRNAs listed in **Table S5** were purchased from Dharmacon. For G9a overexpression, a cDNA encoding the long isoform of G9a was purchased from Dharmacon (clone ID:6822432) and cloned into the pBabe-Puro plasmid using the following primers: forward primer 5’-GTTAGGATCCATGGCGGCGGCGGCGGGAGC-3’, and reverse primer 5’-GTTAGAATTCTTAAGAGTCCTCAGGTGTTG-3’.

Retrovirus was produced by transfecting AmphoPhoenix packaging cells with the retroviral expression construct (National Gene Vector Repository). Lentivirus was produced by transfection of HEK293T cell line with psPAX2 (a gift from Didier Trono, Addgene #12260), pMDG.2 (a gift from Didier Trono, Addgene #12559) and the lentiviral expression construct. Sodium butyrate (1 mM) was added 24 and 48 hours after transfection to boost viral titers. Viral supernatant was collected 48 and 72 hours post-transfection, filtered and used to transduce target cells in the presence of 4 µg/mL polybrene (Sigma). Cells were selected in media containing puromycin (2 µg/mL for primary tumor cells; 4 µg/mL for recurrent tumor cells) and/or blasticidin (5 µg/mL).

### Small molecule screen

Tumor cells from primary #1, primary #2, recurrent #1, and recurrent #2 were plated at 1,500 cells / well on 96-well plates and allowed overnight to attach. Cells were treated with 8 increasing concentrations (vehicle, 10 nM, 30 nM, 100 nM, 300 nM, 1 µM, 3 µM, 10 µM) of each drug in biological triplicate for 60 hours. Each well was normalized to the average of vehicle-treated control wells corresponding to the cell line. Cell viability was determined using the CellTiterGlo kit (Promega) according to the manufacturer’s instructions. The IC_50_ and SEM were determined for each primary and recurrent cohort by combining all 6 biological replicates. The difference in IC_50_ and 95% confidence interval were determined by inputting log[IC_50_] and SEM for each cohort into a web-based tool available at https://www.graphpad.com/quickcalcs/errorProp1/?Format=SEM.

### Cell growth and cell viability assays

Concentration-response curves were determined by plating 1,000 cells from primary and recurrent tumor cell lines in triplicate on a 96-well plate. Cells were allowed to grow overnight prior to 48 hour treatment with increasing concentrations of the following drugs: BIX-01294, UNC0638, BRD4770, Nec-1, GSK’872, and Necrosulfonamide. Cell viability was measured by CellTiterGlo kit. A single IC_50_ with standard error was calculated by nonlinear regression for each cohort and significance between cohorts determined by Student’s unpaired t-test. Concentration-response curves were generated in GraphPad Prism 8 software.

Cell growth kinetics following G9a knockout in primary (#1 and #2) and recurrent (#1 and #2) tumor cells was determined by plating 1,000 cells in triplicate on a 96-well plate. Cell viability was determined using CellTiterGlo on days 1, 3, and 5 post-plating. Relative cell viability was determined by calculating the ratio of luminescence on days 3 and 5 as compared to day 1.

For crystal violet staining, primary or recurrent tumor cells were plated onto 6-well plates (60,000 cells/well) and then treated with 2 µM BIX-01294 for 48 hours. For measurement of cell viability following RIPK3 knockdown, tumor cells were infected with lentivirus expressing one of two shRNAs targeting RIPK3 (shRipk#1 TRCN0000022534, shRIPK3#2 TRCN0000022538) and selected in puromycin. Four days after infection, 50,000 tumor cells were plated onto 10-cm plates and grown for 7-9 days until confluency. Plates were washed with PBS and stained with 0.5% crystal violet for 5 minutes. Excess crystal violet was rinsed with water and plates scanned.

For short-term cell viability following TNF administration, 1,000 cells from each primary (#1, #2, and #3) and recurrent (#1, #2, #3) tumor cell lines were plated in triplicate on 96-well plates and treated for 72 hours with 10 ng/mL TNF (Biolegend) or 0.1% BSA vehicle control. Cell viability was determined using CellTiterGlo. Long-term colony formation assays were performed by plating 1,500 cells on 10-cm plates and treating with 10 ng/mL TNF or vehicle control for 7 days. Plates were washed with PBS and stained with 0.5% crystal violet. Plates were analyzed on Li-Cor Odyssey®. For cell viability assays with necrostatin-1, 1,000 tumor cells from recurrent (#1, #2, and #3) lines were plated in triplicate and treated for 16 hours with 300 nM BIX-01294 alone or in combination with 30 µM Necrostatin-1.

### Immunoblotting and qRT-PCR

Immunoblotting and qRT-PCR was performed as previously described (Mabe et al., 2018). For histone blots, histones were extracted from cell pellets by the protocol found at https://www.abcam.com/protocols/histone-extraction-protocol-for-western-blot. Primary antibodies used for immunoblotting and gene probes used for qRT-PCR are listed in **Table S5**.

### Copy Number Assay

DNA was extracted using the Allprep kit (Qiagen) according to manufacturer instructions and diluted to 5 ng/µL. 7.5 µL master mix was generated for each well in the following ratios: 5 µL genotyping Taqman mix (ThermoFisher Scientific), 0.5 µL of each the Met and Tfrc copy number probes **(Table S5)**, and 1.5 µL of water. 2.5 µL was added to master mix for a total of 10 µL reactions in each well. Samples were run on a CFX384 Real-Time PCR Detection System (BioRad). Met Ct values were normalized to Tfrc using ΔΔCt and compared to normal mouse DNA.

### In-cell western

7,500 tumor cells (Primary #1, Primary #2, Recurrent #1 and Recurrent #2) were plated in duplicate on a black, clear-bottom 96-well plate. Cells were treated for 48 hours with increasing concentrations of BIX-01294 and then fixed with 3.7% formaldehyde for 20 minutes at room temperature, washed with PBS, and permeabilized with 0.1% Triton X-100 in PBS. Cells were blocked for one hour with a 3% goat serum, 1% BSA, 0.1% Triton X-100 solution, prior to overnight incubation of primary antibodies listed in **Table S5**. Secondary antibodies AlexaFluor® (ThermoFisher) 680 and IRDye®800 (Li-Cor) were diluted 1:2000 in PBST and were incubated on cells for one hour. Plates were imaged with a Li-Cor Odyssey® infrared imaging system (Li-Cor Biosciences). Mean fluorescence of each well was analyzed in ImageStudio Lite software (Li-Cor) and the signal ratio determined by dividing the H3K9me2 signal by the Histone H3 signal.

### Flow Cytometry

150,000 tumor cells (Prim #1, Prim #2, Rec #1 and Rec#3) were plated on 6-cm plates and treated for 16 hours with 1 µM BIX-01294 or vehicle control. To test the requirement for necroptosis, recurrent lines #1 or #3 were plated in biological triplicate and treated with vehicle, or 750 nM BIX-01294 with or without 30 µM Necrostatin-1. After treatment, cells were trypsinized and washed with PBS. Cell pellets were resuspended into 100 uL of Annexin-binding buffer (Life Technologies) containing 1:20 Annexin V-AlexaFluor 488 and propidium iodide (Sigma) and incubated for 20 minutes. Samples were diluted with 400 uL of additional Annexin-binding buffer and passed through a 40 μm filter. Samples were excited with a 488 nm laser and 10,000 events were analyzed on a BD FACSCanto II flow cytometer. Data were analyzed using FlowJo software.

### Immune profiling

Orthotopic tumors were manually chopped and digested in a buffer containing: EBSS media (ThermoFisher, #14155063), 1X collagenase (300 Units/mL) / hyaluronidase (100 Units/mL) (Stemcell Technologies, #07912), 2% FBS (Corning, #35010CV), 0.1 mg/mL gentamycin, and 1X penicillin (2000 Units/mL) / streptomycin (2000 Units/mL) (Gibco, #15140-122) for 3 hours at 37°C. Tumors were washed twice with media, and then incubated in a digestion buffer of 5 Units/mL dispase (Stemcell Technologies, #07913) and 0.1 mg/mL DNase I (Worthington, #LS002006). Cell pellets were incubated in ACK lysis buffer for 5 minutes and rinsed twice in FACS buffer (BD Pharmingen, #554656). Tumor cells were strained through a 40 micron filter and resuspended to 1 x 10^7^ cells / mL in FACS staining buffer. 1 x 10^6^ of cells were blocked with 2 µL of CD16/CD32 (Invitrogen, #14-0161-82) for 10 minutes on ice. The cells were then incubated for 30 minutes with antibodies appropriate for cell panels of interest (leukocytes, macrophages, or T-cells) **(Table S5)**. Samples were rinsed twice with staining buffer prior to analysis on BD FACSCanto II flow cytometer and data were analyzed using FlowJo software.

### Orthotopic Recurrence Assays

Nude mice were randomized into cages and maintained on 2 mg/mL doxycycline for two days prior to orthotopic injection. 1 x 10^6^ primary tumors cells (#2) expressing Cas9 and either sgNT, sgG9a#1, or sgG9a#2 were injected bilaterally into the fourth mammary gland of nude mice (n=10 per sgRNA, n=30 mice total). Tumor size was determined by caliper measurement at least three times a week until tumors reached ∼75 mm^3^. Two mice (n=4 tumors) were sacrificed with primary tumors for downstream biochemical analysis. The remaining 8 mice were removed from dox to initiate tumor regression. Mice were palpated at least three times a week to monitor for tumor recurrence (∼75mm^3^). Recurrence-free survival was determined using Kaplan-Meier survival analysis and differences in recurrence were assessed using the Mantel-Cox log-rank test.

### In vivo BIX administration

200,000 recurrent tumor cells (line #3) or 500,000 primary tumor cells (line #2) were injected bilaterally into the inguinal (fourth) mammary gland of FVB or TAN mice, respectively. TAN mice are used as hosts for primary tumor cells because these mice are tolerized to the luciferase protein (data not shown). Mice were randomized to receive vehicle (n=5 mice) or BIX treatment (n=5 mice; 10 mg/kg) by intraperitoneal injections three times a week for two weeks starting one day after tumor cell injections. Tumors size was determined by caliper measurements daily until mouse sacrifice. Mice were sacrificed once tumors reached ∼250 mm^3^. Tumor volume was calculated using the equation ((π x length x width^2^) / 6). Tumor growth curves were compared between treatment groups by repeated-measures, two-way ANOVA (time x treatment) and Sidak’s multiple comparisons test. Tumor AUC was calculated using the equation [(vol_1_ + vol_2_)/2] x (day_2_ - day_1_). Tumor burden was compared between treatment groups by unpaired Student’s t-test.

### ChIP- and RNA-sequencing

Chromatin immunoprecipitation was performed as previously described with antibodies and primers found in **Table S5** (Mabe et al., 2018). Complete RNA- and ChIP-sequencing data is available online using National Center for Biotechnology Information’s Short Read Archive (SRA) under project accession number: PRJNA505839.

### Chip-Seq analysis

Following immunoprecipitation, DNA was quantified using the fluorometric quantitation Qubit 2.0 system (ThermoFisher Scientific) and fragment size confirmed with Agilent Tapestation. DNA libraries were prepared using Kapa BioSystem HyperPrep Library Kit for compatibility with Illumina sequencing. Unique indexes were added to each sample. Resulting libraries were cleaned using SPRI beads, quantified with Qubit 2.0 and Agilent Bioanalyzer, and pooled into equimolar concentrations. Pools were sequenced on Illumina Hiseq 4000 sequencer with 50 bp single reads at a depth of ∼55 million reads per sample. Fastq reads underwent strict quality control processing with the TrimGalore package to remove low quality bases and trim adaptor sequences. Reads passing quality control were mapped to mm10 version of the mouse genome using the Bowtie short read aligner (Langmead et al., 2009). Duplicate reads were filtered and peaks were called with the MACS2 peak-calling algorithm using default parameters, except for H3K27me3 peaks which were called using ‘broad peak’ settings (Zhang et al., 2008). Following sequencing, differential binding analysis was completed with DiffBind software on standard settings (Ross-Innes et al., 2012). Differentially bound sites were annotated with ChIPseeker (Yu et al., 2015) and annotatr (Cavalcante and Sartor, 2017) packages. Enhancer regions were based on Fantom5 classifications. Bam alignment files were converted into bigwig files by binning reads into 100bp segments. Global enrichment plots and heatmap visuals for H3K4me3, H3K9ac, K3K27me3, and RNApol2 at the top 100 most differentially enriched RNApol2 peaks in primary and recurrent cohorts were generated using deepTools2 software (Ramirez et al., 2016). The number of overlapping ChIP-seq peaks with at least one base pair shared was determined with ChIPpeakAnno (Zhu et al., 2010). Publicly available H3K4me1 and H3K27ac ChIP-seq data for mouse embryonic fibroblasts were obtained from ENCODE (via Bing Ren, Ludwig Institute for Cancer Research) and accessed via GEO numbers GSM769028 and GSM1000139, respectively (Consortium, 2012).

### RNA sequencing

RNA was extracted using the RNeasy kit (Qiagen) and concentrations quantified using Qubit 2.0 and Agilent Bioanalyzer. cDNA libraries were prepared with the Kapa stranded mRNA kit. Pooled sample libraries were sequenced on HiSeq 4000 (Illumina) sequencer to 50 bp single reads at ∼25-30 million read depth per sample. Fastq files were assessed for quality control with FastQC software. Raw sequencing reads were then trimmed of low quality base pairs and adapters with Trim Galore! Processed sequencing reads were aligned to the mm10 genome with STAR ultrafast universal RNA-seq aligner (Dobin et al., 2013). The number of transcript counts per gene were counted on the reverse strand with featureCounts software and organized into a count matrix for passing through differential gene expression analysis with DESeq2 software (Love et al., 2014). Of note, recurrent tumor cell line #2 and primary tumor cell line #3 were completed as a separate batch, and were corrected by adding the batch number as an additional variable. PCA analysis was produced using limma and ggplot2 package, and heatmaps were generated with pheatmap package in R (v3.5.2) software. Gene ontology for differential gene expression between primary and recurrent tumor cell lines was determined with clusterProfiler software (Yu et al., 2012). For GSEA analysis, genes were pre-ranked based on topics of interest (i.e. genes differentially expressed in recurrent tumor cell lines or genes upregulated following BIX administration) and input into desktop GSEA software (Broad Institute).

To determine whether the magnitude of gene expression is significantly altered between primary and recurrent tumor cells, the DEseq2 model was designed using Batch + Cell_Line + Condition + Cell_Line:Condition. Significant differences in the magnitude of induction was determined by an adjusted *P* value of <0.05. Recurrent tumor cell line #4 was excluded from these analyses, as we were interested in gene expression alterations specific to recurrent tumor cell lines that were sensitive to G9a inhibition. Significantly altered genes from the analysis were labeled in color on the scatterplot of gene expression changes specific to recurrent cells (y-axis) and primary cells (x-axis) in R software using ggplot2 software.

For heatmap generation of significantly altered genes with BIX treatment, log_2_ gene expression was median-centered within each cohort (i.e. recurrent and primary) to eliminate baseline gene expression differences between primary and recurrent cohorts. Each row was Z-score normalized and input into ‘pheatmap’ software. Genes were clustered according to ‘complete’ method.

### Integrated RNA- and ChIP-sequencing

Genes that were significantly upregulated with BIX in recurrent tumor cell lines #1, #2, and #3 by RNA sequencing (P. adj. <0.05 vs. primary tumor cell lines #1, #2, and #3) were overlapped with gene promoters significantly depleted (P adj. <0.05) of H3K9ac in recurrent tumor cell lines #1, #2, and #3 relative to primary tumor cell lines #1 and #2 by ChIP-sequencing. Dot plot of overlapped data was generated in GraphPad Prism 8 software. Gene ontology of these genes was determined by clusterProfiler package (Yu et al., 2012). This G9a-regulated gene set was input into online software (available at: http://co.bmc.lu.se/gobo/gsa.pl) to determine human distant metastasis-free survival based on a gene set. Settings for this analysis were set for distant metastasis-free survival (DMFS) for all tumors censored at 10 days and divided into 3 quantiles (Ringner et al., 2011). Shown are plots for Luminal B and Her2-enriched tumors. A high G9a signature was determined by low global expression of G9a-regulated genes.

### Statistical Analysis

Western blots show representative results from at least two independent experiments. Gene expression analysis show results from a single representative experiment and are shown as the mean ± the SEM. Two-tailed Student’s t-test was used for statistical analyses between two groups. One-or two-way ANOVA with Sidak’s multiple comparisons test was used to compare statistics across multiple groups. Fisher’s exact test was used to evaluate significance between categorical (i.e. Met amplification) data. Differences in survival were determined by Mantel-Cox log-rank test. A *P* value of <0.05 was considered significant.

## Supplemental Information

Figure S1 includes additional analysis of cell lines features and ChIP-seq analysis. Figure S2 validates on-target effects of G9a inhibition, and shows additional controls from in vivo experiments. Figure S3 provides ChIP-qPCR and gene expression data to support the G9a-regulated gene set reported in the main figures, and presents immune profiling of BIX-treated tumors. Figure S4 includes additional data to support a role of p53 in BIX-mediated cell death. Figure S5 shows the effect of G9a inhibition on a wide panel of human cell lines. Table S1 shows epigenetic drug targets from the screen. Table S2 shows differentially regulated genes following BIX treatment. Table S3 is the integrated G9a-regulated gene set. Table S4 is the curated necroptosis gene set used for GSEA. Table S5 provides a list of antibodies, primer sequences, probes, and additional reagents.

## Acknowledgements

We thank So Young Kim and the Duke Functional Genomics Core for the generation and validation of single guide RNAs targeting G9a and p53; David Corcoran for assistance in the interpretation of ChIP-sequencing data; and Nicolas Devos from the Duke Sequencing and Genomics Technologies Core for acquisition of RNA- and ChIP-sequencing reads. We thank members of the Alvarez lab, including Andrea Walens for assistance with mammary gland injections, Ashley DiMarco for technical assistance with flow cytometry, and Doug Fox for the generation of Cas9-expressing tumor cells. This work was funded by the National Cancer Institute under award numbers R01CA208042 (to JVA), F31CA220851 (to NWM), and F31CA239421 (to RN), as well as the American Cancer Society under award 132556-RSG-18-130-CCG (to JVA) and by startup funds from the Duke Cancer Institute, the Duke University School of Medicine, and the Whitehead Foundation (to JVA).

## Author contributions

NWM, SEW, RN, RCM, BAV generated and analyzed data. RN and RL assisted with animal work. SEW, RCM, and BAV assisted with generation of human cell line treatments. SEW assisted with drug screening and biochemical analysis of p53 knockout. RN performed immune profiling. NWM and JVA wrote and edited the manuscript. JVA supervised all work. All authors reviewed the manuscript.

