## Supplemental Figures and Tables for "G9a Promotes Breast Cancer Recurrence Through Repression of a Pro-inflammatory Program"

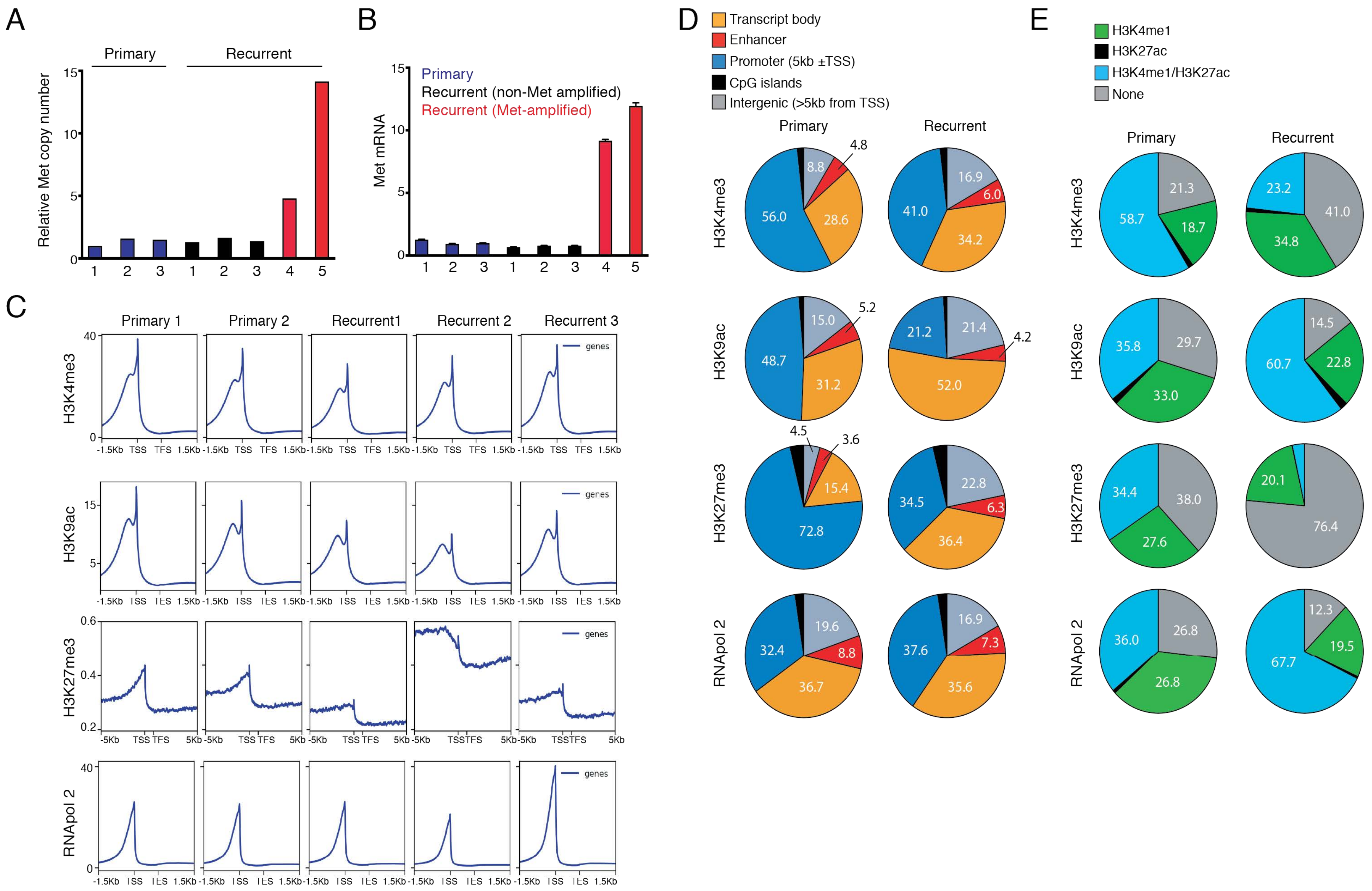
A
Primary
Recurrent
Relative Met copy number
B
Primary
Recurrent (non-Met amplified)
Recurrent (Met-amplified)
Met mRNA
C
Primary 1
Primary 2
Recurrent1
Recurrent 2
Recurrent 3
H3K4me3
H3K9ac
H3K27me3
RNApol 2
genes
-1.5Kb TSS TES 1.5Kb
-5Kb TSS TES 5Kb
D
Transcript body
Enhancer
Promoter (5kb ±TSS)
CpG islands
Intergenic (>5kb from TSS)
E
H3K4me1
H3K27ac
H3K4me1/H3K27ac
None

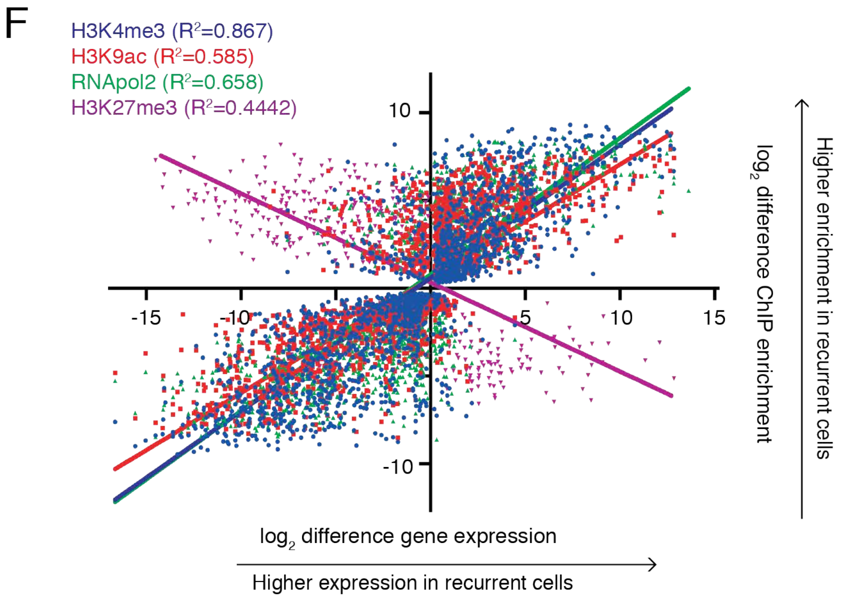


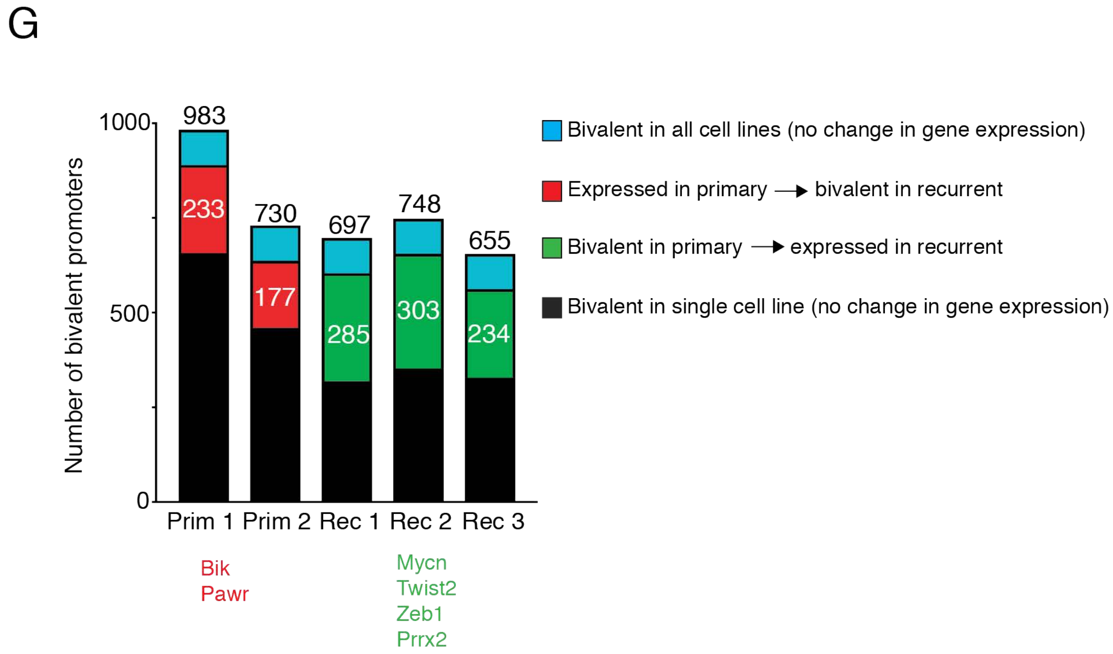


**Figure S1. Tumor recurrence is associated with widespread epigenetic remodeling.**

A) Copy number analysis for Met in primary (n=3) and recurrent (n=5) tumor cell lines. Data are shown relative to recurrent #1.

B) qPCR analysis for Met in primary and recurrent tumor cell lines from panel (A).

C) ChIP-seq enrichment plots for H3K4me3, H3K9ac, H3K27me3 and RNApol2 signal ±1.5 kb (±5.0 kb for H3K27me3) relative to the transcription start site (TSS) and transcription end site (TES) for all genes.

D) Pie charts showing the genomic distribution of unique ChIP-seq peaks in primary and recurrent cells. Promoters are defined as peaks within 5 kb of the TSS.

E) Pie charts showing the percent overlap for H3K4me3, H3K9ac, H3K27me3, and RNApol2 peaks with publically available H3K4me1 and H3K27ac peaks in mouse embryonic fibroblasts.

F) Scatterplot showing correlation of $\log_2$ differential gene expression alterations (x-axis) against $\log_2$ differential ChIP-seq signal (y-axis) for H3K4me3, H3K9ac, H3K27me3 and RNApol 2 epigenetic marks between primary and recurrent tumor cell lines.

G) The absolute number of gene promoters with bivalent histone marks (i.e. both H3K4me3 and H3K27me3) in each cell line. Bivalent promoters are further divided into those that are bivalent in all cell lines (blue), bivalent in a single cell line (black), or bivalent only in primary or recurrent cells and expressed in the opposite cohort (red and green).

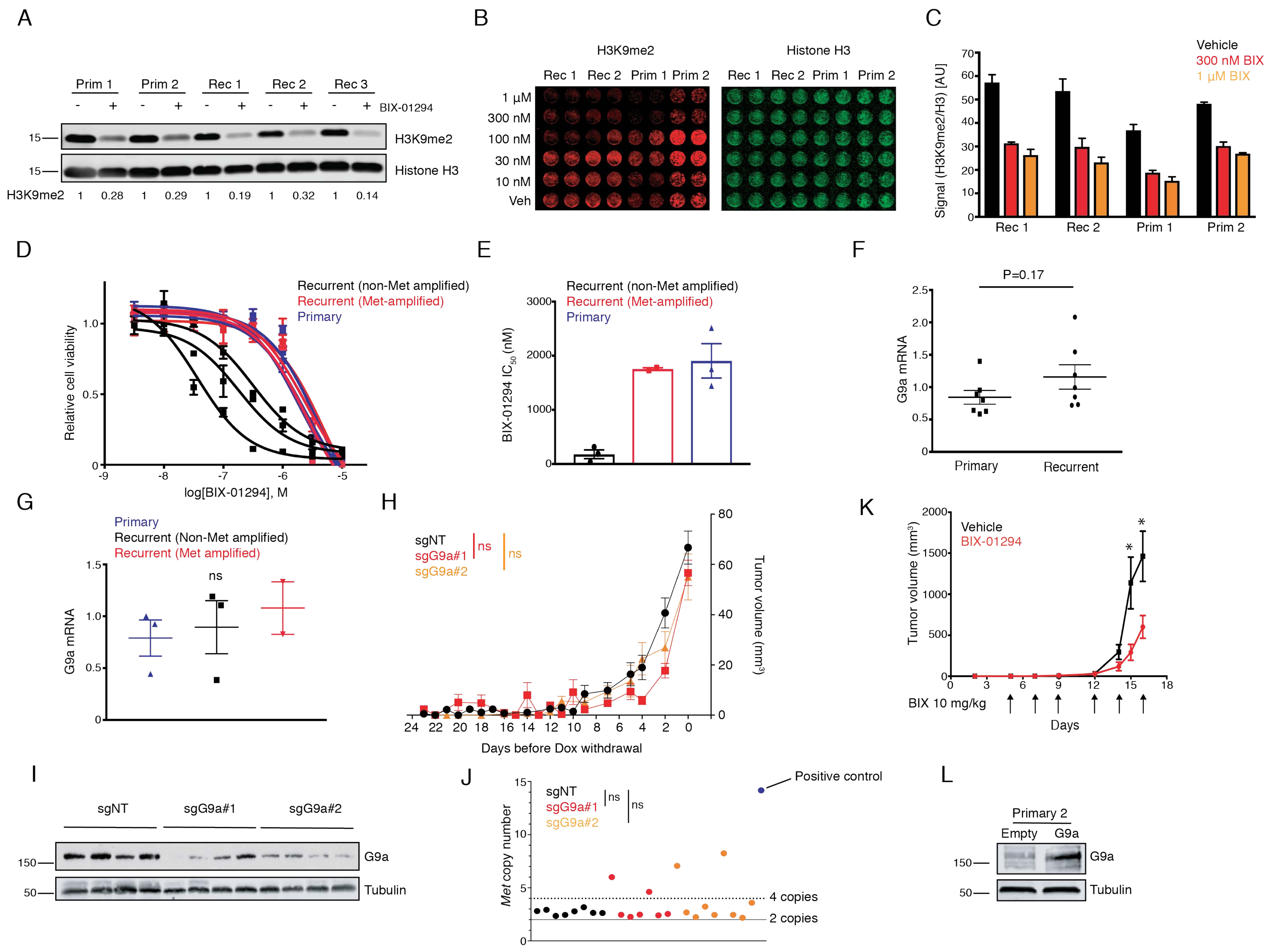
A
Prim 1
Prim 2
Rec 1
Rec 2
Rec 3
BIX-01294
H3K9me2
Histone H3
H3K9me2 1 0.28 1 0.29 1 0.19 1 0.32 1 0.14
B
H3K9me2
Histone H3
Rec 1 Rec 2 Prim 1 Prim 2
1 µM
300 nM
100 nM
30 nM
10 nM
Veh
C
Vehicle
300 nM BIX
1 µM BIX
Signal (H3K9me2/H3) [AU]
Rec 1 Rec 2 Prim 1 Prim 2
D
Recurrent (non-Met amplified)
Recurrent (Met-amplified)
Primary
Relative cell viability
log[BIX-01294], M
E
BIX-01294 IC50 (nM)
F
P=0.17
G9a mRNA
Primary
Recurrent
G
Primary
Recurrent (Non-Met amplified)
Recurrent (Met amplified)
ns
G9a mRNA
H
sgNT
sgG9a#1
sgG9a#2
ns
ns
Tumor volume (mm3)
Days before Dox withdrawal
K
Vehicle
BIX-01294
Tumor volume (mm3)
BIX 10 mg/kg
Days
I
sgNT
sgG9a#1
sgG9a#2
G9a
Tubulin
J
Positive control
sgNT
sgG9a#1
sgG9a#2
ns
ns
Met copy number
4 copies
2 copies
L
Primary 2
Empty
G9a
G9a
Tubulin

**Figure S2. Related to Figures 2 and 3. Recurrent tumors are dependent upon G9a histone methyltransferase activity.**

A) Western blot analysis for H3K9me2 and Histone H3 following treatment with 1 µM BIX-01294 for 6 days. H3K9me2 quantification is shown below each lane and normalized relative to the vehicle control within each cell line.

B) In-cell western showing H3K9me2 (red) and Histone H3 (green) protein in primary and recurrent tumor cells treated with increasing concentrations of BIX-01294 for 48 hours.

C) The ratio of H3K9me2 to H3 mean fluorescence intensity for 300 nM and 1 µM values from (A). Significance was determined by one-way ANOVA and Tukey's post-hoc test.

D) Concentration response curves for primary, non-Met amplified recurrent, and Met-amplified recurrent tumor cells treated with increasing concentrations of BIX-01294.

E) Bar plot showing individual $IC_{50}$ values calculated from (D).

F) qPCR analysis showing G9a transcripts for a panel of independent primary (n=7) and recurrent (n=7) tumors. Significance was determined by Student's unpaired t-test with P-value indicated.

G) qPCR analysis showing G9a transcripts for a panel of primary (n=3) and recurrent (n=5) tumor cell lines. Significance was determined by one-way ANOVA and Tukey's posthoc test.

H) Tumor growth curves for sgNT, sgG9a#1, or sgG9a#2-expressing primary tumors (n=20 tumors / cohort) in nude mice. Statistical significance was determined by repeated-measures 2-way ANOVA (time x sgRNA). Significance values are shown for the last day between sgNT and each sgRNA as determined by one-way ANOVA and Tukey's posthoc test.

I) Western blot analysis showing G9a expression in representative control or G9a-knockout recurrent tumors.

J) Copy-number analysis for the *Met* gene in sgNT (n=8), sgG9a#1 (n=7), or sgG9a#2 (n=9) recurrent tumors. Met values were normalized relative to tail DNA. The solid and dashed lines indicate 2 copies and 4 copies of Met, respectively. Tumors with greater than 4 copies of Met were considered amplified. Significance was determined by Fisher's exact test.

K) Mean tumor growth curves for orthotopic recurrent tumors in nude mice treated with vehicle or BIX-01294 (10 mg/kg). Arrows indicate drug treatments.

L) Western blot analysis showing G9a expression in primary tumor cell line #2 expressing either an empty vector or a cDNA encoding the long isoform of G9a.

Error bars denote mean ± SEM. ns= not significant

A

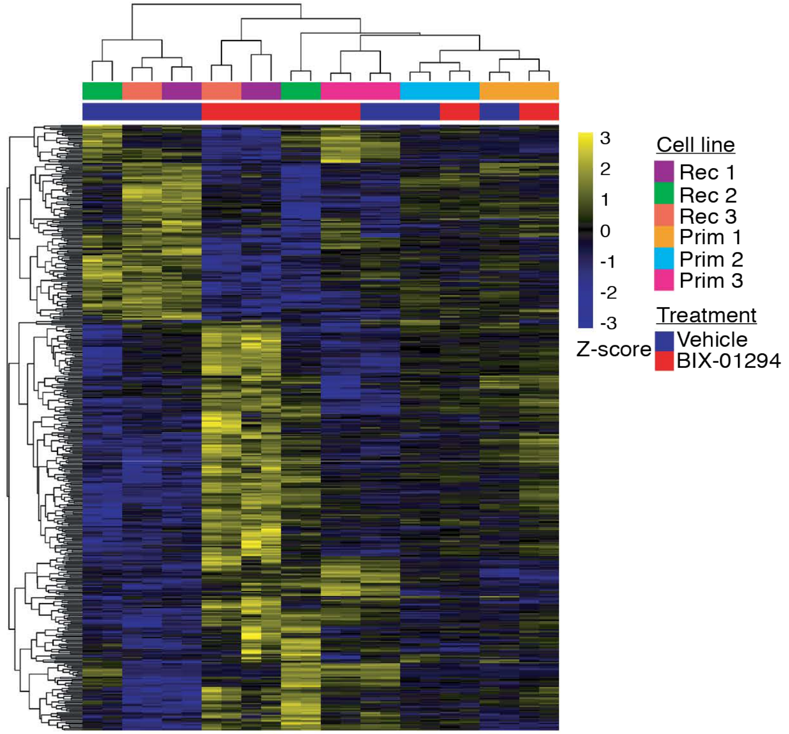
3
2
1
0
-1
-2
-3
Z-score
Cell line
Rec 1
Rec 2
Rec 3
Prim 1
Prim 2
Prim 3
Treatment
Vehicle
BIX-01294

B

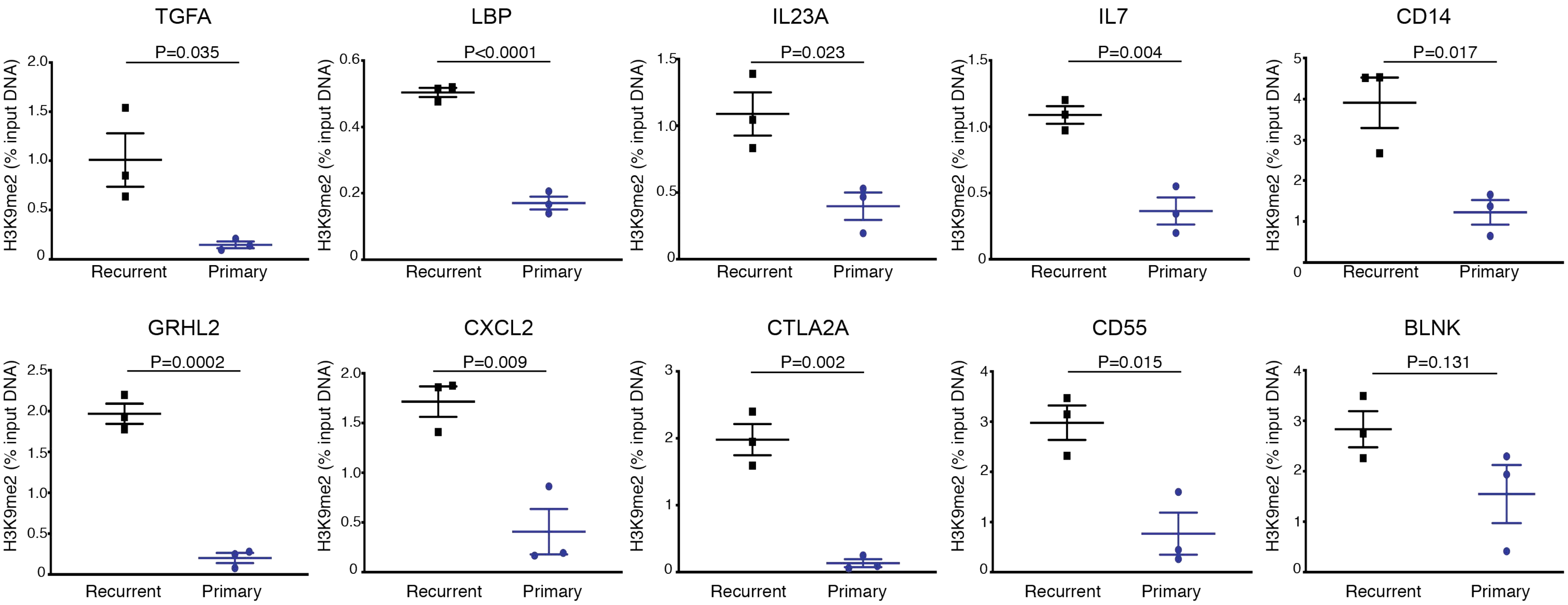
TGFA
P=0.035
LBP
P<0.0001
IL23A
P=0.023
IL7
P=0.004
CD14
P=0.017
GRHL2
P=0.0002
CXCL2
P=0.009
CTLA2A
P=0.002
CD55
P=0.015
BLNK
P=0.131
H3K9me2 (% input DNA)
Recurrent
Primary

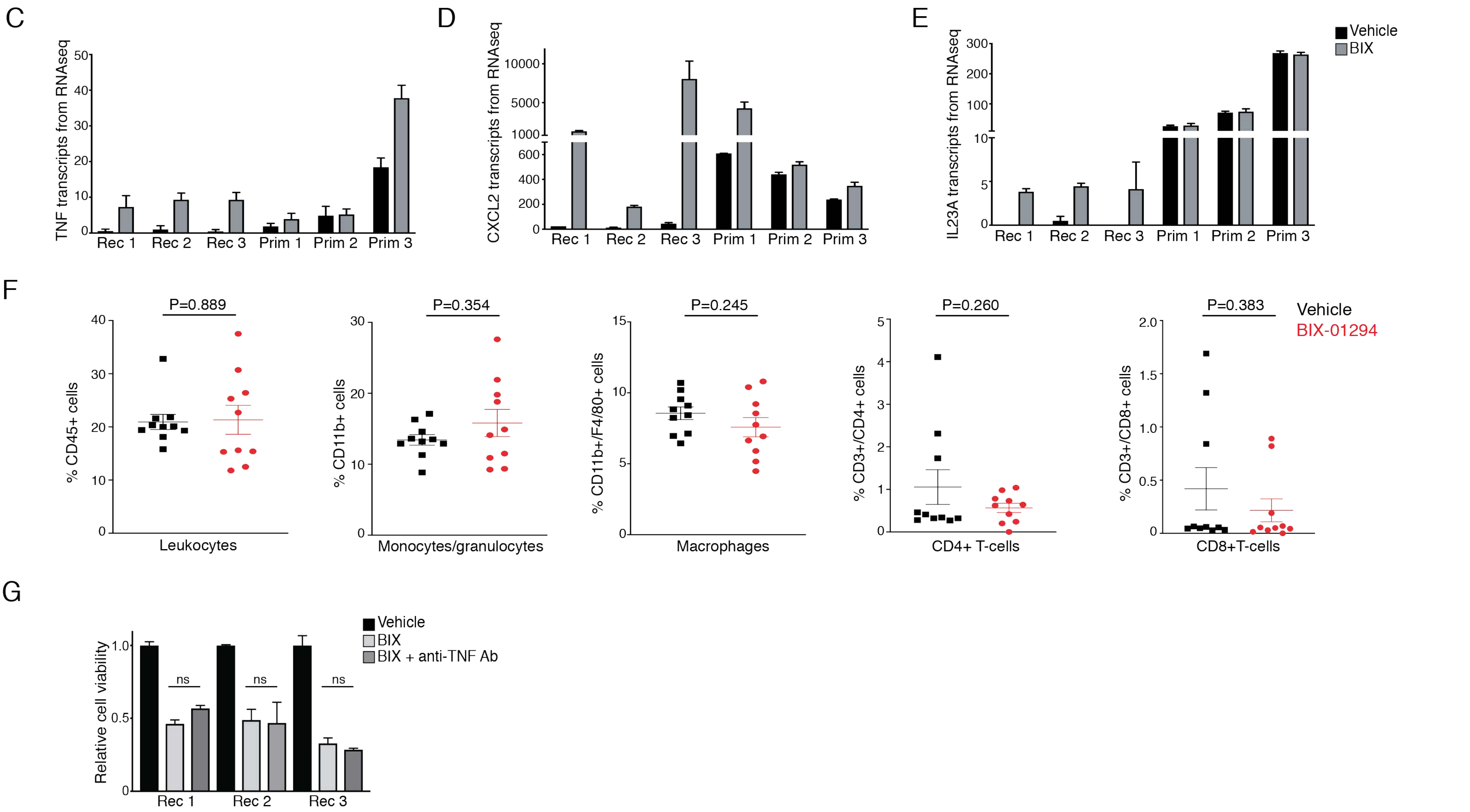


**Figure S3. Related to Figures 4 and 5: Integrated epigenetic and transcriptional analysis of G9a-regulated genes in recurrent tumors.**

A) Heatmap showing unsupervised hierarchical clustering of median-centered, Z-score normalized gene expression changes following BIX treatment.

B) ChIP-qPCR showing H3K9me2 enrichment at the promoters of indicated genes from the G9a-regulated gene set. Each point represents an independent cell line. Significance was determined by Student's unpaired t-test and *P*-values are indicated.

C-E) Normalized RNA-seq transcript counts for TNF (C), CXCL2 (D), and IL23A (E) in recurrent and primary tumor cell lines following BIX treatment.

F) Immune cell profiling by flow cytometry showing the abundance of leukocytes (CD45+), monocytes/granulocytes (CD11b+), macrophages (CD11b+/F4/80+), and T-cells (CD3+/CD4+ or CD3+/CD8+) in recurrent tumors treated with vehicle (n=10) or BIX-01294 (n=10). Significance was determined by Student's unpaired t-test and *P*-values are indicated.

G) Cell viability in recurrent cells treated with 300 nM BIX-01294 for 3 days in the presence or absence of an anti-TNF neutralizing antibody. Significance was determined by one-way ANOVA and Tukey's post-hoc test.

Error bars denote mean ± SEM. ns= not significant

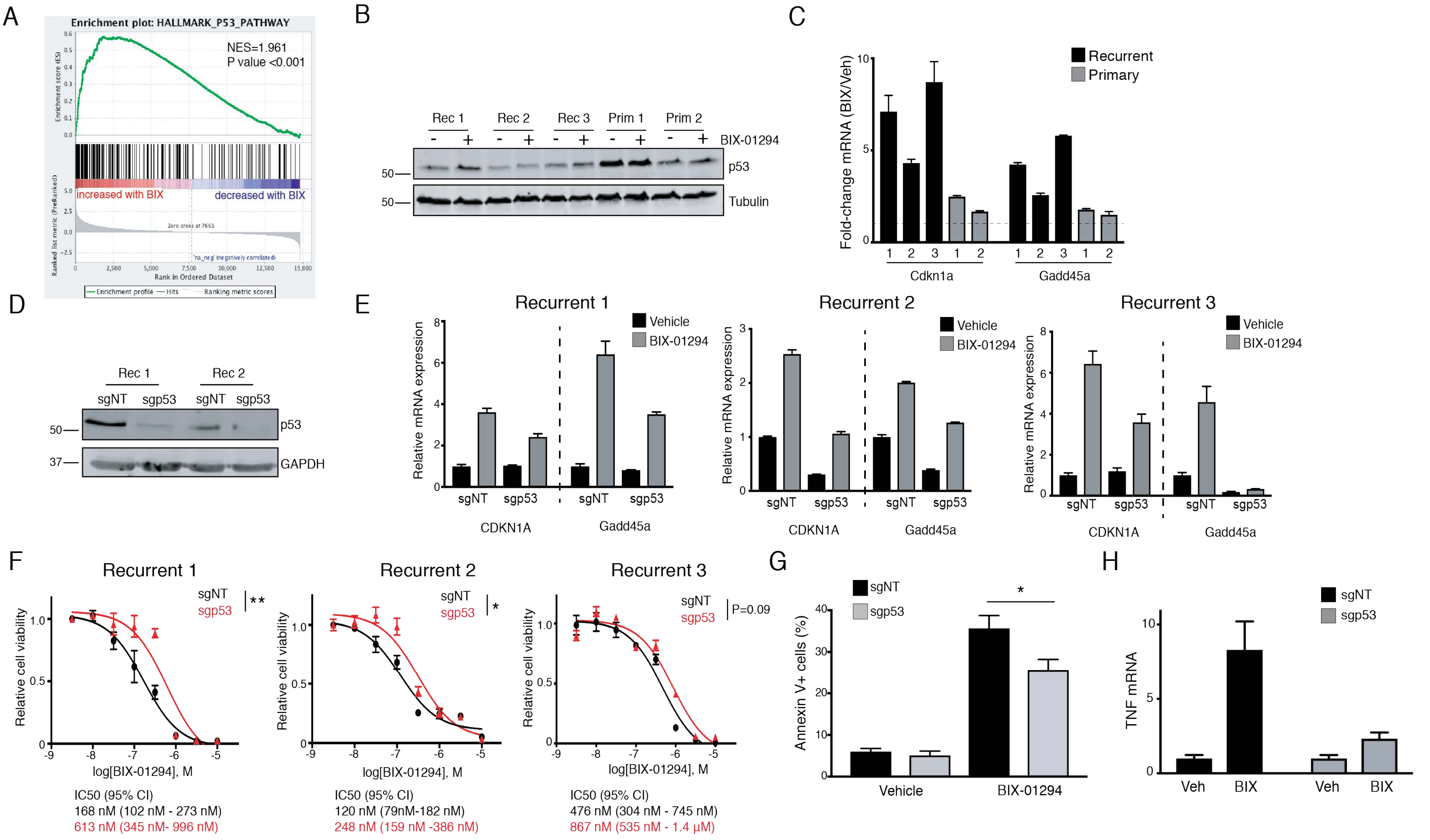


**Figure S4. G9a leads to induction of p53 targets and p53-dependent cell death.**

A) GSEA plot showing enrichment for a p53 signature in recurrent tumor cells following G9a inhibition. *P*-value and normalized enrichment score are shown.

B) Western blot analysis showing p53 expression in primary and recurrent tumor cell lines following treatment with vehicle or 1 µM BIX.

C) qPCR analysis of p21 (Cdkn1a) and Gadd45a expression following BIX treatment in primary and recurrent tumor cells. Data are expressed as fold-change relative to vehicle within each cell line.

D) Western blot showing p53 expression in recurrent tumor cell lines expressing Cas9 nuclease and a single guide RNA targeting p53.

E) qPCR analysis of Cdkn1a and Gadd45a expression in control and p53 knockout recurrent tumor cells following 1 µM BIX administration.

F) Concentration response curves for control or p53 knockout recurrent tumor cells treated with increasing concentrations of BIX-01294. $IC_{50}$ values were calculated for each cohort by non-linear regression and significance was evaluated by Student's unpaired t-test.

G) Annexin V staining of recurrent tumor cell line #1 expressing sgNT (n=3) or sgp53 (n=3) after 24 hours treatment with 750 nM BIX-01294.

H) qPCR analysis for TNF expression in control or p53-knockout recurrent tumor cells (#3) treated with 1 µM BIX-01294.

Error bars denote mean ± SEM. *P<0.05, **P<0.01

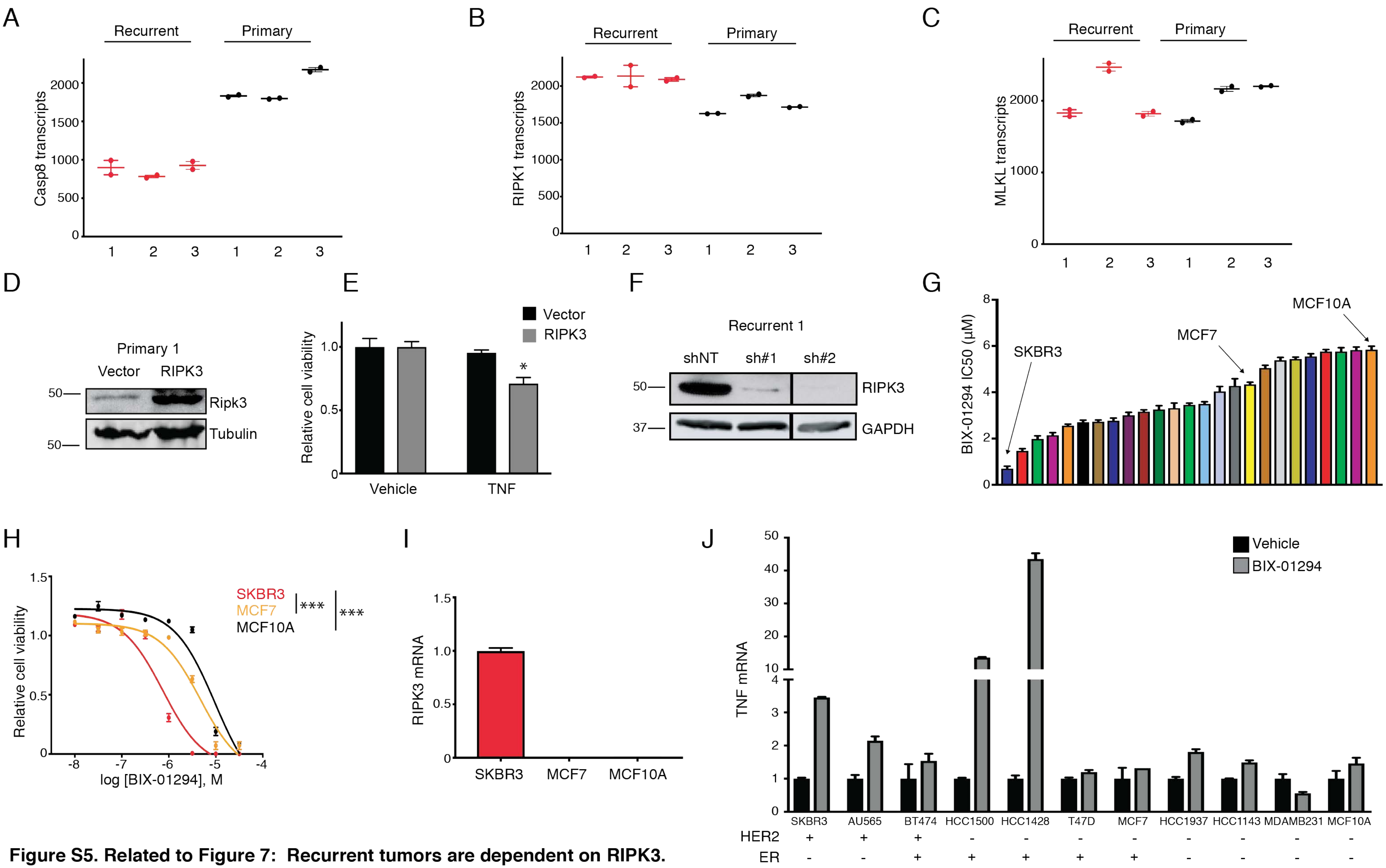


**Figure S5. Related to Figure 7: Recurrent tumors are dependent on RIPK3.**

A-C) Normalized RNA transcript counts for Casp8 (A), RIPK1 (B), and MLKL (C) in primary and recurrent tumor cell lines as determined by RNA-seq.

D) Western blot analysis showing RIPK3 expression in primary line #1 transduced with retrorvirus expressing a cDNA encoding RIPK3.

E) Cell viability following 72 hour treatment with 10 ng/mL TNF in empty vector or RIPK3-overexpressing primary cells. Significance was determined by one-way ANOVA and Tukey's posthoc test.

F) Western blot analysis showing RIPK3 expression in recurrent line #1 following knockdown with one of two shRNAs targeting RIPK3.

G) Concentration response curves with BIX-01294 were determined for 24 human breast cancer cell lines and MCF10A immortalized mammary cells. Waterfall plots indicate $IC_{50}$ values for each cell line and standard error. Cell lines SKBR3, MCF7 and MCF10A are indicated.

H) Concentration response curves with BIX-01294 for SKBR3, MCF7 and MCF10A human cell lines.

I) qPCR analysis for RIPK3 expression in SKBR3, MCF7 and MCF10A cell lines.

J) qPCR analysis for TNF expression in a panel of human breast cancer cell lines following 48 hours vehicle or 4 µM BIX-01294 administration.

Error bars denote mean ± SEM. ****P*<0.001. N.D. denotes not detected.

**Table S1: List of epigenetic drug targets**

| Target |
|---|
| DNA methyltransferases |
| Protein arginine demethylase 1,3,4 |
| BAZ2A/B |
| BRD4 |
| P300/CREB |
| SMARCA |
| SIRT2 |
| JMJD2 |
| PHD1/2 |
| O-6-methylguanine DNA methyltransferase |
| LSD1 |
| BRD2/4 |
| PARP |
| DOT1L |
| Pan-JMJC |
| SET7/9 |
| EZH2 |
| L3MBTL3 |
| SUV39H1 |
| AURKA |
| G9a |
| HDAC4 |
| HDAC5 |
| HDAC1/2 |

**Table S3: G9a-regulated gene set in recurrent cells**

2010300C02Rik
4930523C07Rik
A330023F24Rik
A530046M15Rik
ACE
ACER2
ADARB1
ADGRG3
AKAP9
ALDOC
ALS2
AMPD3
AP5Z1
ARAP2
AREG
ARHGAP27
ARHGEF3
atg9a
ATP6V1F
ATP8B1
B3GNT3
BC031181
BHLHE41
BLNK
BMT2
BTBD3
BTC
BTN1A1
C77080
Car2
Car7
CARD14
CD14
CD55
CDCP1
CDKN2AIP
cds1
CELSR1
CFLAR

CHKA
chrnb1
CISH
CLCN3
CLDN23
CNKSR1
CNPY4
COBL
COL18A1
CORO6
CREM
CSF2
Ctla2a
CTSD
CTTNBP2NL
CXCL2
CYP51
CYSTM1
DACT2
DCTN4
DDIT4
DDIT4L
DDT
DLG1
DLL1
DNAJB5
DOK7
DTNA
DUSP13
dusp28
DUSP6
DYRK3
ECHDC1
EIF2AK3
ENPP5
EPB41L5
EPHB3
EPHB4
ERLIN2
ESCO1

ESRP2
ETV3
EXPH5
F11R
FAAH
FAM167A
fam83h
FAM84A
FBXL20
FBXO7
FBXW9
FDPS
FERMT1
FGD4
FGD6
FGF1
FRRS1
FUCA2
GABARAP
gbp7
GCA
gcnt2
GDPD1
GIPC2
GLMP
Gm19510
Gm38426
GNA14
GNPDA2
GPR137
GPX4
GRAMD1C
GRHL2
GstT3
H2-Q1
H2-Q2
H2-T23
HES6
HK1
HOXA5

HSD17B7
HSDL1
igsf11
IL23A
IL24
IL7
IRX3
ITGA2
JUNB
jund
KANK1
KBTBD2
KCNH1
KCNN4
KCNQ1
KIF17
KISS1R
KLC3
LAMC2
LBP
LIF
LLGL1
LLGL2
LMTK2
LONRF1
LPIN3
LRBA
LSMEM1
LSS
Ly6a
MALT1
MAP3K9
MCTP2
MEGF10
MFAP3L
MFSD6
MICALL1
MKL2
MNT
MOCOS

MON1B
MPP4
MRGPRE
MTMR3
MVD
MXD1
MYBL1
NAA40
NABP1
NAIF1
NARF
Nat8f4
NCK2
NDRG4
NEDD4L
NEU1
NHLRC3
NRARP
NUDT12
Oasl1
OCIAD2
OPTN
OSGIN1
PACSIN3
PAFAH1B3
PAPLN
PARD6B
PCDH1
Pea15a
PELI1
PEX19
PEX26
PHACTR2
PHLDA1
PHLDA2
PHLPP1
PICK1
PID1
PIKFYVE
PIP5K1B

PLCG2
PLEKHA4
PLEKHA7
PLEKHH1
PLET1
PLIN3
PLK3
pmvk
PPFIBP2
PRDM1
PRKCZ
PRMT2
PROSER2
PRRG4
PRSS22
PSTPIP1
PSTPIP2
PTPN12
PTPRJ
PTTG1IP
RAB11FIP1
RAB15
RAB21
RABGEF1
RC3H1
REPIN1
rfxap
RHBDF2
RHOU
rin2
RNF11
RNF146
RNF152
RPTOR
rubcn
RUNX1
SAMD10
SCLY
SDCBP
SEC11C

SEC14L2
SH3BP2
SHF
SHISA4
SKIL
SLC12A6
SLC12A9
SLC25A48
SLC27A1
SLC35A3
SLC35E3
SLC35F1
SLC40A1
SLC45A3
SLC4A11
SLC52A3
SLC5A3
SLC6A15
SLC7A4
SLC8B1
SLC9A3R1
Slfn2
SNX16
SOCS2
SORL1
SORT1
SOX9
SP140
SPAG9
SPIDR
SPINT1
SPINT2
SQSTM1
SRGAP1
ST14
st6gal1
STAM2
STAP2
STX7
STYK1

SULT2B1
SUN1
SUSD1
SYNRG
TBC1D30
TBC1D8
TEC
TECPR1
TFAP2A
TFAP2C
TFRC
TGFA
TIMM22
TJP2
TLE6
TLR3
TMC7
TMCC1
TMCO3
TMEM128
TMEM140
TMEM14A
TMEM184A
TMEM40
TMEM62
TMEM71
TNF
TNFAIP3
TNFRSF11A
TNIK
TNIP2
TNS4
TOB1
TRAF4
TRAFD1
TRIM7
Trp53inp1
TSPAN14
TTC9
UBE2Q1

UNC13B
urah
Utp14b
VAMP1
VPS26A
VPS37B
VPS50
WDR26
WDR81
WFS1
WNT9A
WWC1
zbed4
ZBTB38
ZBTB7B
ZFAND2A
Zfp212
Zfp418
Zfp655
Zfp664
Zfp68
Zfp7
Zfp750

**Table S4: Curated necroptosis gene set for GSEA**

ABCE1
ABCF1
ADAM26A
ADO
ARFRP1
ATP6AP1
ATP6AP2
BBC3
BIRC2
BIRC3
BMF
BOK
C030046E11RIK
CAD
CASP8
CBFB
CCDC101
CFLAR
COG1
COG3
COG4
COG6
COMMD4
CREBL2
CSF2
CWC15
CXCL1
CXCL2
CYLD
DDI2
DDX6
DEFB1
DMXL1
DNML1
Dpysl4
ecd
EIF5B
EPHX1
FADD

FAS
Foxi1
Fth1
Galnt5
GGNBP2
GLUD1
GLUL
GPR152
GRB2
GRID2
HERC4
HOXA3
HSPBAP1
INSM2
IPMK
Jph3
Kcnip1
LSG1
Mag
MAP3K7
MAPK1
Mapk8
Mapk9
MCTS1
MED12
MEPCE
METTL3
MLKL
Mrcl
N4BP1
NFE2L1
NFKBIA
NMD3
NRBP1
NUDCD2
NXT1
OGDH
OXSR1
parp2
PCBP1

PELI1
PGAM5
plrg1
PPIF
PPP2CA
PPP6C
PSMD6
PTBP1
PVR
PYGL
RAB25
RAD23B
RAI14
rasa4
RBCK1
RGP1
RIPK1
ripk3
RNK31
RPL10
RPL37
S100A7A
SEC23IP
SFRS2
SFT2D2
SIRT2
SLC1A5
SLC25A4
SLC7A11
SMG7
SMG8
SPATA2
SRCAP
STRAP
stx1a
TADA1
TEX10
TLR3
TMED9
TMEM107

TMEM57
TNF
TNFAIP81L
TNFRSF1A
TRAF1
TRMT11
Trp53
TRPM7
TXB3
Txn14b
UBE2K
UFC1
UPF2
URB1
USP9X
VPS29
WNK1
XRN1
YTHDC1
ZBP1
ZEB2
ZSWIM8

**Table S5: Reagents**

| Antibodies | | | | |
|---|---|---|---|---|
| Antibody | Source | Application | Catalog | Dilution |
| H3K9me2 | Cell Signaling | WB | #4658 | 1:1000 |
| Histone 3 | Cell Signaling | In-cell western ; WB | #3638 | 1:500 (ICW), 1:1000 (WB) |
| G9a | Cell Signaling | WB | #3306 | 1:1000 |
| GAPDH | Sigma-Aldrich | WB | #8795 | 1:2000 |
| Tubulin | Cell Signaling | WB | #3873 | 1:2000 |
| p53 | Cell Signaling | WB | #2524 | 1:1000 |
| Cleaved PARP | Cell Signaling | WB | #9544 | 1:1000 |
| Cleaved Caspase | Cell Signaling | WB | #9661 | 1:1000 |
| MLKL | Cell Signaling | WB | #37705 | 1:1000 |
| p-MLKL(S345) | Abcam | WB | #196436 | 1:1000 |
| RIPK3 | Santa Cruz | WB | #374639 | 1:500 |
| H3K9me2 | Abcam | ChIP-qPCR, In-cell western | #1220 | 5 μg ; 1:500 (ICW) |
| H3K9ac | Abcam | ChIP-seq | #4441 | 5 μg |
| H3K4me3 | Abcam | ChIP-seq | #8580 | 5 μg |
| H3K27me3 | Abcam | ChIP-seq | #6002 | 5 μg |
| RNA pol II CTD repeat YSPTSPS | Abcam | ChIP-seq | #817 | 5 μg |
| Mouse IgG isotype control | Abcam | ChIP | #18392 | 5 μg |
| Rabbit IgG isotype control | Abcam | ChIP | #171870 | 5 μg |
| AlexaFluor® 680 anti-rabbit | ThermoFisher | WB | #A21076 | 1:5000 |
| IRDye® 800 anti-mouse | Li-Cor | WB | #926-32210 | 1:5000 |
| Goat anti-rabbit IgG (H + L)-HRP conjugate | Biorad | WB | #1706515 | 1:5000 |
| CD11-PE | BD Biosciences | Flow Cytometry | #561689 clone M1/70 | 1:50 |
| F4/80-AF647 | BD Biosciences | Flow Cytometry | #565853 clone T452341 | 1:50 |
| CD3e-PE | BD Biosciences | Flow Cytometry | #561824 clone 1452C11 | 1:100 |
| CD8a-APC | BD Biosciences | Flow Cytometry | #561093 clone 53- | 1:200 |

| | | | | |
|---|---|---|---|---|
| | | | 6.7 | |
| CD4-APCCy7 | BD Biosciences | Flow Cytometry | #561830 clone GK1.5 | 1:100 |
| CD45 - APC | BD Biosciences | Flow Cytometry | #561018 clone 30-F11 | 1:200 |
| ChIP-qPCR primers | | | | |
| TNFα Forward | 5'-GCCTTTATAGCCCTTGGGGA-3' | | | |
| TNFα Reverse | 5'-GAGACAGAGGTGTAGGGCCA-3' | | | |
| BLNK Forward | 5'-GGGGCTTTCTGCTCATTTGC-3' | | | |
| BLNK Reverse | 5'-ATGACCACTGCTGTGACCTC-3' | | | |
| IL23A Forward | 5'-TGGGATTCCCCTCCCTACAT-3' | | | |
| IL23A Reverse | 5'-TTTCCCCTGGAACTGAAGCG-3' | | | |
| CD55 Forward | 5'-AGCTACCCACACTCCCAACT-3' | | | |
| CD55 Reverse | 5'-ATTTCTCAGCTCCCTCTGGCT-3' | | | |
| GRHL2 Forward | 5'-GTCTCTCCCATCACCTCTACCC-3' | | | |
| GRHL2 Reverse | 5'-TGCGAAGTTTACCTGAGTGGA-3' | | | |
| TGFA Forward | 5'-CCCTAGCACAGGTGACTGGAT-3' | | | |
| TGFA Reverse | 5'-GTCCCCTGCGTAGTAGCC-3' | | | |
| CTLA2 Forward | 5'-GTCACAAGCCAGCTTCGCTA-3' | | | |
| CTLA2 Reverse | 5'-GACATCCTGTGAGTCTTGGCA-3' | | | |
| CXCL2 Forward | 5'-CCAGACTCCAGCCACACTTC-3' | | | |
| CXCL2 Reverse | 5'-CGAATCCGAATCCCACCTGT-3' | | | |
| LBP Forward | 5'-TATAGGCTGGCCAAGGAGGC-3' | | | |
| LBP Reverse | 5'-GCCACATACCCGTATAGCAGA-3' | | | |
| IL7 Forward | 5'-TGATTAAGAGGGTCACGCCC-3' | | | |
| IL7 Reverse | 5'-TCCTACCTGAGCAGGTGCAT-3' | | | |
| CD14 Forward | 5'-GCATTAGGGCATTCTGTGCC-3' | | | |
| CD14 Reverse | 5'-CCCTTTTCCTGTACGCACCA-3' | | | |
| Taqman gene expression probes | | | | |
| Cdkn1a | ThermoFisher Scientific | Mm00432448 | | |
| Ehmt2 | ThermoFisher Scientific | Mm01132261 | | |
| Tnf (mouse) | ThermoFisher Scientific | Mm00443258 | | |
| Ripk3 (mouse) | ThermoFisher Scientific | Mm00444947 | | |
| Actb | ThermoFisher Scientific | Mm02619580 | | |
| 18s | ThermoFisher Scientific | 4332641 | | |
| Gadd45a | ThermoFisher Scientific | Mm00432802 | | |
| Met | ThermoFisher Scientific | Mm01156972 | | |
| TNF (human) | ThermoFisher Scientific | Hs00174128 | | |
| RIPK3(human) | ThermoFisher Scientific | Hs01011175 | | |
| CRISPR-Cas9 sgRNAs | | | | |
| sgNT | 5'-CCCGATCCCCTACCTAGCCG-3' | | | |
| sgG9a#1 | 5'-CGGCAGGCTCCAAGGAGTCG-3' | | | |
| sgG9a#2 | 5'-ACAGGCACCCCCCTTGCTGG-3' | | | |

| | | |
|---|---|---|
| sgp53 | 5’-GAAGTCACAGCACATGACGG-3’ | |
| Copy number assay probes | | |
| Met | ThermoFisher Scientific | Mm00565151_cn |
| Tnfr | ThermoFisher Scientific | 4458366 |
| Small hairpin RNAs | | |
| shScrambled | Addgene | #1864 |
| shRipk3#1 | Dharmacon | TRCN0000022534 |
| shRipk3#2 | Dharmacon | TRCN0000022538 |
